# Deep Proteoform Sequencing with Top-Down Direct Mass Technology

**DOI:** 10.64898/2026.05.29.728917

**Authors:** Kenneth R. Durbin, Taojunfeng Su, Ryan T. Fellers, John P. McGee, Nickolas P. Fisher, Michael A.R. Hollas, Jared O. Kafader, Neil L. Kelleher

## Abstract

Individual Ion Mass Spectrometry (I^2^MS) using Direct Mass Technology mode on an Orbitrap mass spectrometer (DMTm) increases sensitivity, resolution, and mass range for protein analysis. Here, we present an end-to-end workflow for deep proteoform sequencing using top-down mass spectrometry with DMTm. By assigning the charge of individual fragment ions and converting spectra from the *m/z* to the mass domain, DMTm resolves overlapping isotopic distributions that have limited conventional top-down mass spectrometry. Across different fragmentation modes on Orbitrap mass spectrometers, top-down DMTm significantly outperformed conventional top-down mass spectrometry methods. For a glycosylated 50.8 kDa antibody heavy chain, sequence coverage was greatly increased, from 27.5% to 83.3%, in 10 minutes of acquisition using a single fragmentation mode. Coverage of the middle 350 residues improved from 0% to >95%, demonstrating near-complete coverage of the difficult-to-characterize internal region of a large protein. The fragmentation patterns of DMTm were found to be complementary to conventional top-down, with higher internal coverage for DMTm and higher terminal coverage for conventional. Accordingly, aggregation of the data from the two modes further increased heavy chain sequence coverage to 90.2%. A new software platform, Proteoform Studio, provided optimized ion processing for improved sequence coverage and enabled real-time experimental monitoring as individual ions were accumulated. The platform automatically integrates conventional and DMTm data to provide the most comprehensive sequence coverage possible. Together, these advances enable substantially deeper proteoform sequencing and establish a straightforward, complete top-down DMTm workflow to confidently define proteoforms in biological systems and biotherapeutic development.

## Introduction

Proteins exist in a wide variety of molecular forms, known as proteoforms, which arise from genetic variations, alternative splicing, and post-translational modifications (PTMs).^1, 2^ This molecular diversity plays a critical role in maintaining protein structure and binding affinity, thereby influencing cellular functions and biological processes. As a result, proteoforms are gaining more attention as promising biomarker candidates. Recent translational studies have demonstrated strong associations between specific proteoforms or overall proteoform landscapes and certain diseases.^3^ In the therapeutic space, protein-based modalities must have well-defined proteoforms (i.e., product or critical quality attributes) to maintain efficacy, stability, and safety.^4^ Comprehensive characterization of proteoforms, including precise determination of sequence and site-specific modifications, is therefore essential for understanding disease mechanisms and guiding the development of targeted therapeutics.^5^

Mass spectrometry technology is a widely used analytical technique for the characterization of proteins because of its speed, accurate mass measurement, and high compatibility for analyzing a wide range of analytes, from peptides to protein complexes.^6, 7^ Top-down mass spectrometry is a powerful technology that directly measures and characterizes intact proteins, without the need for enzymatic digestion during sample preparation.^8, 9^ Compared to bottom-up, top-down mass spectrometry avoids the protein inference problem and provides accurate information on sequence variations, as well as the occupancy and the crosstalk of PTMs.^10^ However, conventional mass spectrometry technologies, also called ensemble measurements, have diminished analytical power for intact proteins over ∼50 kDa.^11^ A similar issue appears for fragment ions from top-down, with fragment ions larger than ∼25 kDa generally not being detected.^12^ One of the main issues for large ion detection is the significant signal overlap that occurs in the mass-to-charge (*m/z*) domain, which reduces the number of clean isotopic distributions that can be algorithmically detected with high accuracy and sensitivity. As a result, only a relatively small fraction of fragments can be determined, and in general, these fragments are primarily small, higher abundance ions near the terminal ends of the protein. This limited fragment coverage can lead to the loss of critical sequence information, such as complementarity-determining region (CDR) 3 in antibodies or key phosphorylation sites close to the middle of proteins.^13-15^ Consequently, these challenges have significantly hindered the adoption of traditional TD-MS in protein characterization.

To alleviate the inherent limitations in TD-MS, different tandem MS activation methods have been introduced to fragment intact proteins, aiming at maximizing the fragmentation efficiency to increase sequence coverage. Higher-energy collisional dissociation (HCD)^16^, electron-transfer dissociation (ETD)^17^, and ultraviolet photodissociation (UVPD)^18^ are three commonly used activation methods for intact protein fragmentation. HCD is broadly compatible with different mass detectors and mainly produces *b*- and *y*-ions through vibrational excitation. It is a robust method but can induce neutral losses on fragments and generate abundant internal fragments that limit terminal sequence coverage and complicate spectral analysis.^19^ In contrast, ETD is considered a mild activation method that preserves labile PTMs and mainly produces *c*- and *z*-ions, often also producing higher sequence coverage.^20^ It is noteworthy that ETD efficiency decreases for precursors with low charge states due to insufficient charge density to drive fragmentation, causing no dissociation after electron transfer. UVPD employs excimer lasers (e.g., 193 nm) to induce extensive backbone fragmentation, generating up to nine types of ions; thus, this method often produces the highest sequence coverage.^21^ However, UVPD demands precise ion control to reach optimal fragmentation, and the diversity of produced fragment ions elevates complexity in data interpretation. One way to maximize the sequence coverage is to combine fragment data derived from different activation methods with a variety of parameter settings.^22^ By combining fragment ions derived from different fragmentation types such as HCD, ETD, EThcD, and UVPD as well as PTCR, these strategies can achieve >90% and >80% sequence coverage of light chain (Lc) and heavy chain (Hc) for antibodies, respectively.^23, 24^ Notably, even with multiple fragmentation types, top-down sequence coverage is significantly lower than peptide mapping, and key regions, particularly in the inner region of the protein, may not be fully sequenced. Other limitations, including the availability of multiple activation methods, ion assignment accuracy, and the amount of instrument acquisition time per analyte, must be considered when implementing a multi-modal fragmentation strategy.

The recent development of charge detection mass spectrometry on Orbitrap systems with Individual Ion Mass Spectrometry (first described as I^2^MS and then commercialized as Direct Mass Technology mode or DMTm) has transformed the capabilities of MS for proteoform mass measurements.^25^ DMTm directly determines charge states of individual ions using the Selective Temporal Overview of Resonant Ions (STORI) plot analysis, bypassing reliance on deconvolution of ions in the *m/z* domain.^26^ This unique charge determination grants two major advantages for the analysis of large biomolecules. First, it can determine the masses of large proteins and complexes in the megadalton range (e.g., adeno-associated viruses).^27^ Second, it significantly improves resolution of analytes in highly complex samples. In contrast, conventional measurements will generally produce convoluted signals forming a broad spectral hump in the *m/z* domain, making proteoform identification unachievable.^25^

Top-down protein characterization also benefits from direct mass measurement.^28^ Previous works have stated that DMTm enhances the sensitivity of fragment ion measurements for top-down MS. Application of top-down DMTm to endogenous proteins has shown that DMTm has high accuracy and sensitivity in identifying post-translational modification (PTM) sites and distinguishing splicing variants.^15, 29^ Here, we demonstrate a complete workflow for deep proteoform sequencing using top-down DMTm. We applied insights from the development of the workflow to produce nearly complete sequence coverage of reduced antibody, showcasing significant strides over what was previously possible for large biomolecule characterization by top-down mass spectrometry.

## Methods

### Protein Sample Preparation

Three recombinant standard proteins, myoglobin, carbonic anhydrase, and pyruvate kinase (Thermo Fisher Scientific, Waltham, MA), and two monoclonal antibodies, SILuMab (Sigma-Aldrich, St. Louis, MO) and NISTmAb, were reconstituted in 70:30 water:acetonitrile with 0.2% formic acid to a final concentration of 1 µM. SILuMab antibody was first incubated with Dynabeads Protein G (Invitrogen) in 1× Tris-buffered saline (TBS) buffer for 4 h at room temperature. Beads were washed three times with 1× TBS and incubated with PNGase-F (Promega, Madison, WI) with 50 mM ammonium bicarbonate overnight at 37 °C. The antibody was then eluted in 100 mM glycine at pH 2 from beads, and the supernatant was subjected to 50 mM TCEP for 1 h at room temperature to be reduced to light chain (Lc) and heavy chain (Hc).

The reduced subunits were desalted with a 10 kDa molecular-weight-cutoff filter (Sartorius AG, Göttingen, Germany) and resuspended in the denaturing buffer. Samples were then centrifuged at 4 °C to remove magnetic beads before mass spectrometry analysis.

### Protein Direct Infusion

Each protein sample except NISTmAb was directly infused at ∼1.5 µl/min via a laser pulled (Sutter P-2000, Novato, CA) fused silica capillary emitter tip (300 µm O.D. × 100 µm I.D.) and ionized using an in-house built electrospray ionization (ESI) source. The spray voltage was set to 2 kV, the capillary inlet was heated to 300 °C, the RF lens was set to 60%, the source-induced dissociation value was set to 15 eV, and the ion routing multipole pressure was set to 2 mTorr. NISTmAb subunits were analyzed via a heated ESI (HESI) source connected to a syringe pump flowing at 3 µL/min. All DMTm and ensemble data were collected as above with spray voltage set to 3 kV.

### Mass Spectrometry Analysis

A Thermo Scientific Orbitrap Eclipse mass spectrometer was used for the analysis of the three standard proteins, a Thermo Scientific Orbitrap Ascend mass spectrometer was used to analyze SILuMab, and a Thermo Scientific Orbitrap Tribrid Apex mass spectrometer with an IR laser was used to analyze NISTmAb (Thermo Fisher Scientific, San Jose, CA). All three instruments were set to developer mode with DMTm enabled. Ensemble MS^1^ data were acquired independently for each protein sample to assess the charge distributions and determine the charge states and *m/z* windows for DMTm data acquisition using 7500 resolving power (**Figure S1**). For the three standard proteins, each selected *m/z* peak was fragmented using HCD, ETD, and UVPD, respectively. The best observed fragmentation condition with ETD was applied to the Lc and Hc of SILuMab. Ensemble MS^2^ data were collected for SILuMab subunits with ETD. Precursor ions with charge states +22 and +57 were selected for fragmentation of the Lc and Hc, respectively, using a 2 *m/z* isolation window to avoid co-isolation. All DMTm data collected on the Orbitrap Eclipse mass spectrometer used a resolution of 500,000, automatic gain control (AGC) and eFT were disabled, µscans were set to 1, and injection time (IT) was manually tuned for each acquisition to maximize the number of individual ion events in each scan. In general, individual ions were collected in 50 min with a collection window of 500-2000 *m/z* for each protein sample.

For SILuMab Lc and Hc DMTm data collection, multiple ETD parameters with 3, 6, and 9 ms were collected for 60 min each. ETD with 35 ms reaction time was used for SILuMab ensemble data and spectra were collected for 25 min.

For NISTmAb, DMTm data were collected for 10 min for each subunit and ensemble data were collected for ∼10 seconds using resolution of 240,000, an Automated Ion Control target of 1000, and a window of 400-2000 *m/z*. For NISTmAb Lc analyzed with IR-EThcD in DMTm, ETD reaction time was set to 3 ms, reagent target was set to 5e5, IR laser power was kept at 8%, and supplemental HCD activation was set to 10% without normalization to charge state. For IR-EThcD ensemble analysis of NISTmAb Lc, ETD reaction time was set to 10 ms, while the rest of the parameters were kept the same as the DMTm analysis. For NISTmAb Hc analyzed by IR-EThcD in DMTm, ETD reaction time was set to 1 ms with 10% charge state normalized supplemental HCD activation; reagent target and IR laser power were kept the same between Lc and Hc. The same parameters as for the Lc ensemble were used for NISTmAb Hc IR-EThcD ensemble analysis with the exception of charge state normalized supplemental HCD set to 10%.

### DMTm and Ensemble Data Processing and Analysis

The charge assignment for individual ions was performed with Proteoform Studio v1.0 (Thermo Fisher Scientific). Fragment ion annotation and manual inspection were performed using the Targeted Top-Down workflow in Proteoform Studio. To manually validate each annotation, a fragment tolerance of 2 ppm, a sub-tolerance of 2 ppm, a minimum signal-to-noise (S/N) ratio of 3, and a minimum ion fitter score of 0.5 were applied with the isotopic fitter in the workflow to retain as many presumptive annotations as possible. Annotations were classified as false identifications if they met any of the following criteria: (1) absence of the top three most abundant isotopic peaks, (2) more than half of the isotopic peaks deviated from the theoretical isotopic distribution, or (3) more than half of the peaks were shared with other isotopic distributions. Proteoform Studio was used to integrate DMTm spectral matches with ensemble spectral matches.

## Results and Discussion

### Converting spectral congestion into productive fragment information

The primary advantage of charge detection mass spectrometry lies in its ability to resolve the charge state of individual ions and then convert all ions into a true mass spectrum. The mass conversion relies on accurate assignments of charge states, which can be difficult given the inherent imprecision in exactly what charge each ion may have. Direct Mass Technology on Orbitrap mass spectrometers (DMTm) enables charge detection at scale by allowing thousands of ions to be analyzed in each spectrum. We use a ‘voting’ algorithm that utilizes neighboring isotopic and charge state ion data across all scans to more accurately assign the charge state of each ion. The thousands to millions of ions collected in each acquisition can therefore be analyzed together to refine charge assignments. Through the voting process, ions with slope values that previously would have been assigned the incorrect charge state are corrected. We optimized the slope assignment and voting algorithm specifically for top-down fragment ions and were able to retain substantially more ions, which led to increased sequence coverage (**Figures 1** and **S2-S3**).

**Figure 1.**
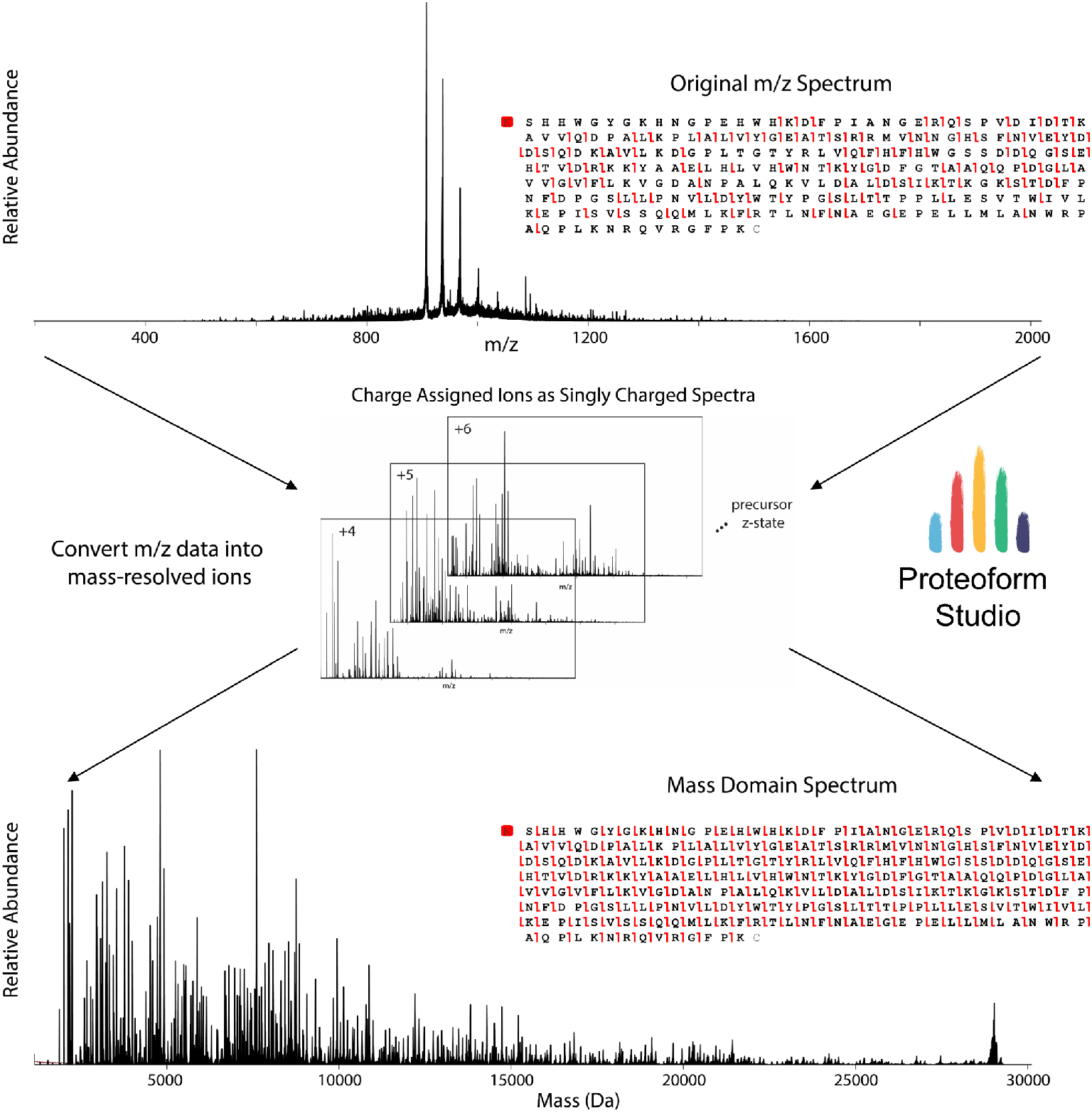
Charge detection improves protein sequence coverage. The majority of the original averaged spectrum from composite data (top) is composed of charge-reduced precursor ions with a high baseline that limited fragmentation coverage. By collecting top-down fragmentation data in Direct Mass Technology mode, each individual ion can be charge assigned using the Proteoform Studio analysis platform (middle), yielding ions with charge states from +4 to the charge state of the isolated precursor. All charge assigned ions can then be combined together and directly converted into a mass domain spectrum (bottom). Theoretical isotopic distributions can be fit to singly charged mass spectra to yield fragment maps. The charge state insets are *m/z* spectra composed of only ions assigned to a specific charge state.

We visualized the *m/z*-domain distribution of ions detected from DMTm spectra by converting the ions of matched carbonic anhydrase fragment ions back into *m/z* values. We found the mean ion counts for each *m/z* bin in the densest regions were approximately 5 to 15 ions per *m/z* bin depending on fragmentation type (**Figure 2A**). These data highlight the major advantage of DMTm; because the data can be analyzed directly in the mass domain, overlaps in *m/z* space are no longer a hindrance to fragment ion detection and assignment. The ion distribution overlaps can be seen across a wide swath of the *m/z* spectrum and form a normal distribution from 500 to 2000 *m/z* range. In **Figure 2B**, a five-*m/z* slice of a representative *m/z* spectrum shows two matching fragment ions. In comparison, we matched 16 ions in the mass spectrum composed of the same group of ions, which illustrates how a reduction in ion overlap substantially impacts the number of fragment ions that can be matched.

**Figure 2.**
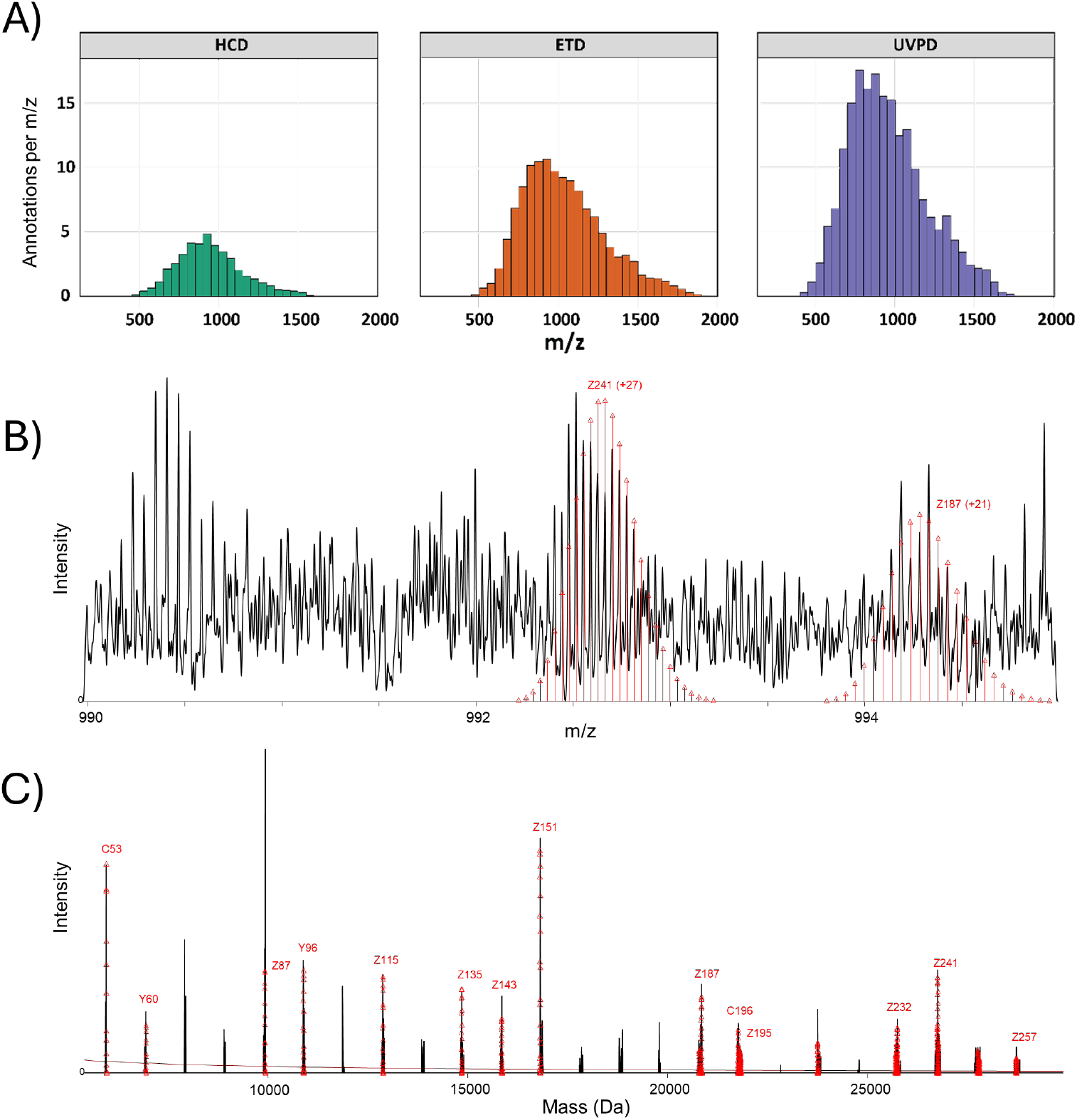
Disentangling ion distributions in top-down spectra. Top-down fragmentation produces *m/z* spectra with overlapping ions in the same m/z space. (A) Histograms were created with 1 *m/z* width bins to find the mean number of overlapping ions per *m/z* for 50 *m/z* windows across the spectrum. (B) A representative section of a 5 *m/z* region from a DMTm-ETD spectrum of carbonic anhydrase is shown with two matching fragment ions annotated. (C) Following conversion of the same *m/z* ions into the mass domain, 16 fragment ions were matched.

### Robust proteoform sequencing with different fragmentation techniques

To further assess the advantages and limitations of DMTm for top-down protein sequencing, we next examined fragmentation patterns and sequence coverages of a 29 kDa protein, carbonic anhydrase, across three different fragmentation methods. After mapping fragment ions in the mass domain, distinct patterns emerged that were dependent on the fragmentation technique.

HCD and UVPD showed clear fragmentation preferences for specific amino acids, with certain fragment ions generated at significantly higher levels compared to other ions. Additionally, both HCD and UVPD predominantly generated smaller, abundant fragment ions, with a significant reduction in the number and abundance of larger fragment ions (i.e., >10 kDa). Fragmentation using ETD generated fragment ions that were more evenly distributed across the mass domain. Although the larger ions had lower intensities than the smaller ions in ETD, their isotopic distributions were well-resolved for confident fragment mapping. Furthermore, ETD preserved abundant precursor ions, offering intact mass information even without MS^1^ data, while the precursor signal was nearly undetectable in HCD and UVPD spectra.

Overall, performing top-down DMTm with ETD achieved the highest sequence coverage, with 93.4% coverage when looking exclusively for *c-* and *z*-type ions from carbonic anhydrase (**Figure 3A**). In contrast, HCD yielded 52.7% sequence coverage, punctuated by preferred fragmentation channels and distinct sequence tag runs (**Figure 3B**). Furthermore, a high number of fragment ions produced by HCD corresponded to internal fragment ions (i.e., ions generated via multiple fragmentation events, see **Figure S4**). UVPD yielded 86.4% sequence coverage, which was comparable to the ETD sequence coverage (**Figure 3C**). However, protein fragmentation with UVPD produces more ion types (all ion types (a, b, c, x, y, z) can be generated and were considered), making fragment annotation more difficult due to many overlapping isotopic distributions. The major complementary sequence information offered by HCD and UVPD to that of ETD was cleavage at the N-terminus of proline residues, which is not readily generated by ETD, suggesting that these methods can provide some amount of complementary information.

**Figure 3.**
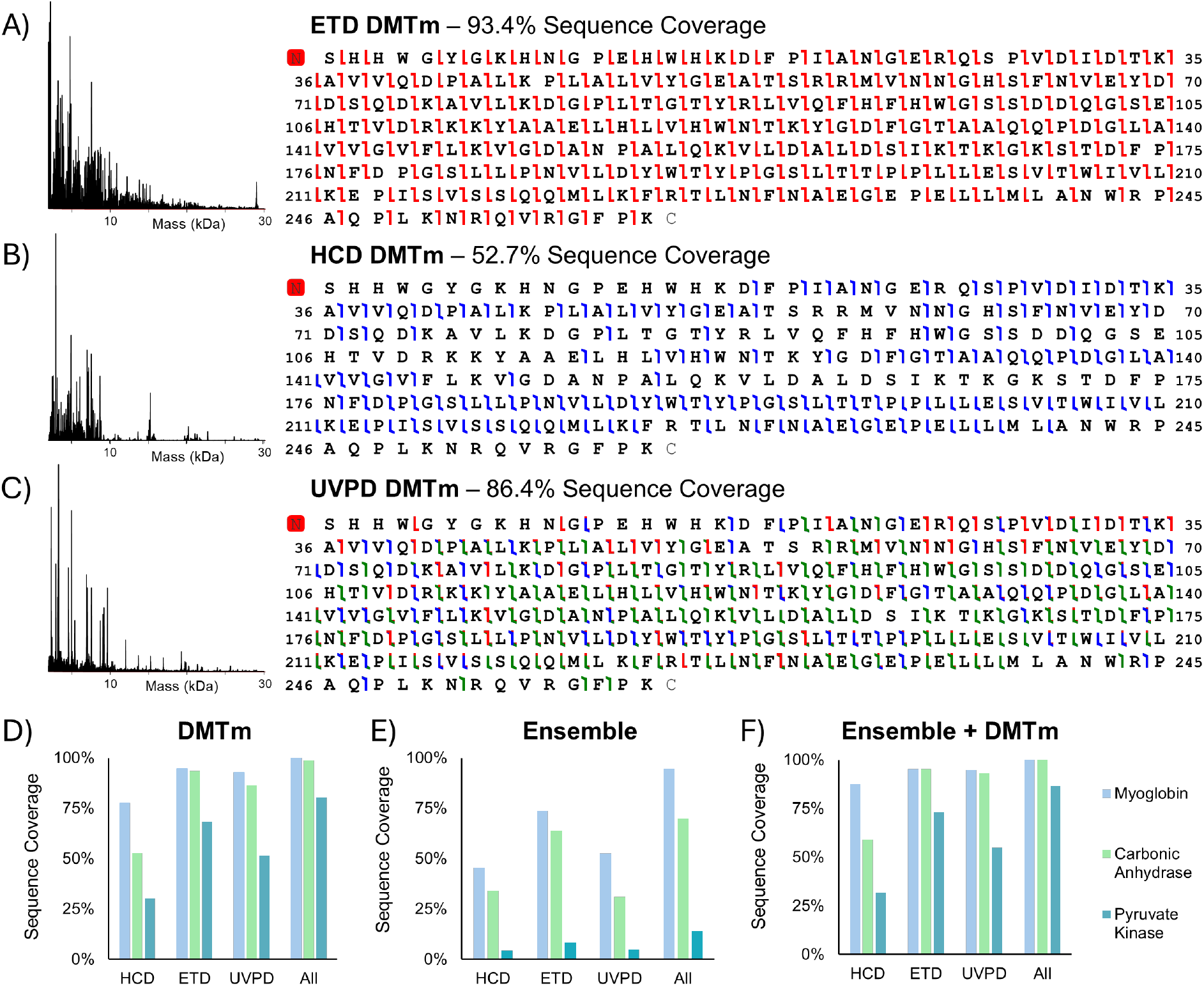
Fragmentation results using top-down DMTm. The 29 kDa protein, carbonic anhydrase, was fragmented with (A) ETD, (B) HCD, and (C) UVPD. (D) Sequence coverage for myoglobin, carbonic anhydrase, and pyruvate kinase is plotted for DMTm only (left), ensemble only (middle), and DMTm combined with ensemble (right).

We next compared the sequence coverage for two different-sized standard proteins using both DMTm and ensemble acquisition modes with HCD, ETD, and UVPD (**Figure 3D**). First, using top-down DMTm, ETD-only fragmentation of myoglobin achieved 94.7% sequence coverage, while UVPD and HCD yielded 92.8% and 77.6% sequence coverage, respectively. The achieved sequence coverage with top-down DMTm was significantly lower for pyruvate kinase, likely due to some combination of large size, sequence, incomplete unfolding, and a high number of disulfides that could make uniform disulfide bond reduction a challenge. ETD still outperformed the other two methods with a sequence coverage of 68.4% compared to 51.6% for UVPD and 30.2% for HCD. Looking closer at the spectra for pyruvate kinase, many larger fragment ions were seen with strong signal-to-noise and clear isotopic distributions (**Figure S5**). When combining DMTm data from all three fragmentation methods, the total sequence coverages were improved, achieving 100%, 98.8%, and 80.3% for myoglobin, carbonic anhydrase, and pyruvate kinase, respectively, indicating each method generated complementary sequence information (**Figure S6**). Across all proteins and fragmentation methods, sequence coverage obtained using ensemble measurements was significantly lower than those using DMTm (**Figure 3D-E**). This disparity became clearer when fragmenting large proteins like pyruvate kinase, where ETD ensemble measurements resulted in only 8.1% sequence coverage for pyruvate kinase compared to 68.4% using ETD with DMTm. In addition, the combined ensemble data from all three fragmentation methods never reached full sequence coverage for any of the proteins studied.

Overall, ETD in DMTm yielded consistently higher sequence coverage than other available techniques, even when considering only two types of fragment ions. Therefore, ETD could be considered as a preferred starting point for deep characterization of intact proteins with DMTm.

### DMTm and ensemble are complementary modes for top-down MS

DMTm has a limitation in that ions with a charge state less than +4 are not detected. We therefore also analyzed the m/z spectra that are collected along with the DMTm data (referred to as composite spectra) to see if they could be mined for additional, terminal ions with lower charge states (see SI section for more details). However, these composite data did not reveal significant gains to sequence coverage. Conversely, comparing ensemble to DMTm revealed distinct and potentially complementary differences, with ensemble fragmentation patterns presenting strong coverage of the terminal ends of the proteins and DMTm fragmentation patterns showing significantly more coverage in the middle portions of the proteins (**Figures S7-S8**). We therefore combined ensemble data with DMTm data to see if the combination would increase the total achievable sequence coverage. Furthermore, this combination did alleviate much of the limitation of DMTm toward detecting ions with lower charge states. Following result combination, we observed significant gains in overall sequence coverage (**Figures 3F** and **S9**). While some proteins may have complete or nearly complete coverage with DMTm, for those proteins with lower DMTm coverages, these results suggest ensemble measurements can offer valuable sequence information (especially near the termini) that may not be accessible via DMTm alone.

### Optimization for accuracy of automated fragment ion assignment

Top-down protein characterization often requires manual validation to remove inaccurate annotations, particularly false positives that can lead to overestimation of sequence coverage and lowered confidence in PTM site localization. However, such validation is tedious and requires expertise from the validator, ultimately limiting the deployment of top-down protein characterization. Although software tools can help automate this process, false positive fragment ion assignments are still commonly observed. With an expert user performing manual validation, more permissive first-pass analysis settings can generally be used, as any data can be filtered during post-processing when the analyst validates the data. Analysis settings for fragment matching can be modified to achieve comparable results as to those achieved with expert manual validation.^23^ To gauge the impact of analysis settings on the expected decoy matches with DMTm data and move toward automating the analysis process while still producing high-fidelity results, we inspected parameter thresholds for ppm tolerance, the isotopic fitter score, and the S/N ratio. Our goal was to produce the lowest number of expected decoy matches while still retaining high sequence coverage. Randomized and shuffled sequences were used for decoy trials. The settings used for the manually validated spectra produced average decoy matches for both shuffled and random sequences corresponding to 16.9% and 11.3%, respectively, of the total fragment matches observed with the base proteoform (**Figure S11A**). The average sequence coverages were 27.2% and 18.7% for shuffled and random decoy sequences. As expected, the shuffled sequences had higher homology with the base proteoform sequence and thus displayed a higher average number of matching fragment ions and sequence coverages for all data points.

Making parameters more conservative led to fold-change reductions in the number of decoy matches seen (**Figure 4A-B**). The fragmentation type also played a significant role in the number of decoy matches observed. With UVPD, more ion types are generated, so the data density and probability of a match are substantially higher than for HCD or ETD where only two ion types are being matched (**Figure 4C**). Comparing ETD with HCD, much higher decoy numbers are seen for ETD, as the spectra are denser due to the more complete and even fragmentation across the spectrum. As expected, the number of residues in the sequence also influenced the average number of decoy matches (**Figure S11B**).

**Figure 4.**
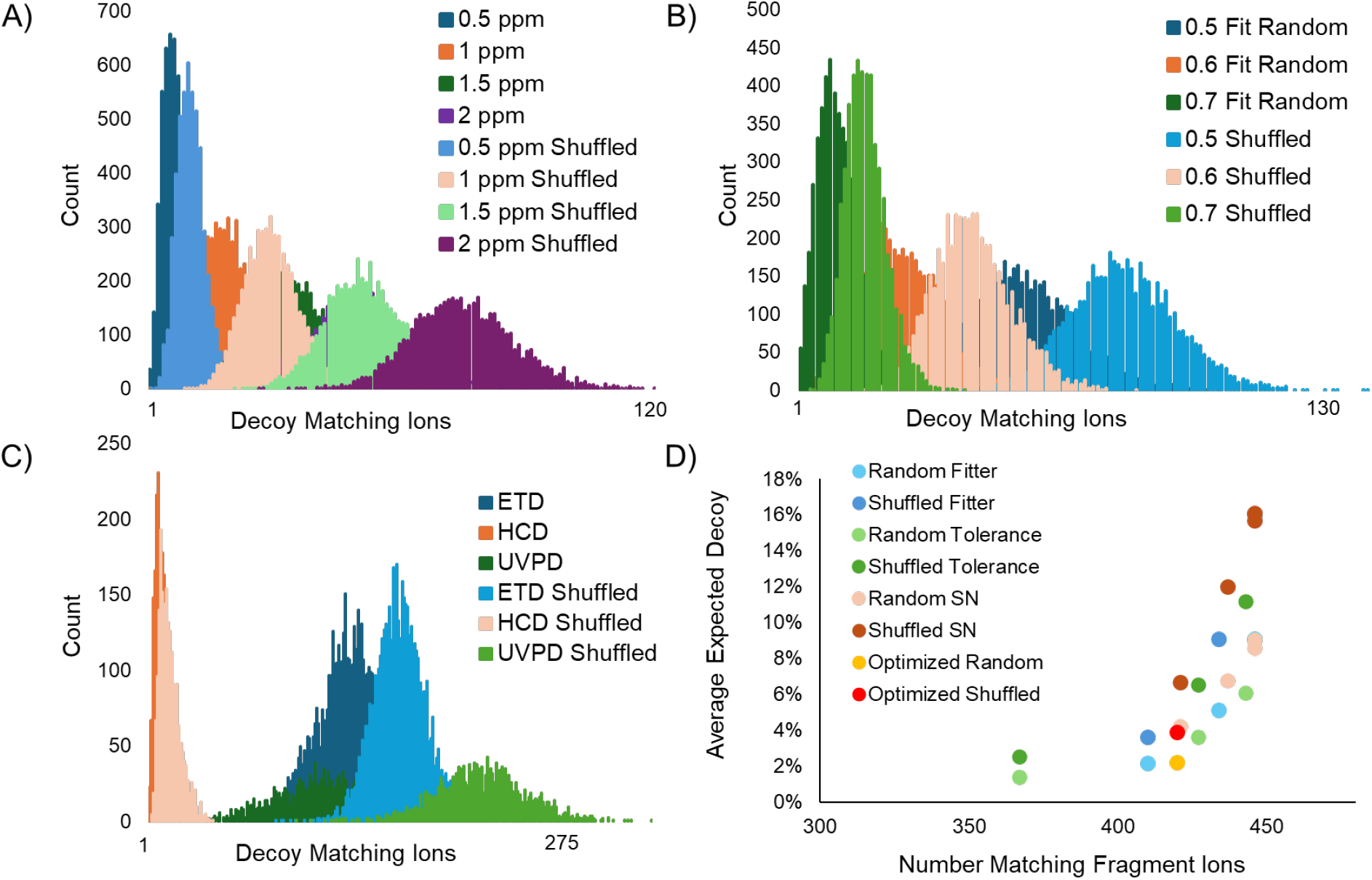
Spectrum-level decoy assessment of DMTm data. Shuffled sequences of carbonic anhydrase and randomized sequences of the same length were analyzed using the isotopic fitting routine against the experimental data. Decoy sequence trials were conducted with (A) different ppm fragment tolerances and (B) different isotopic fit scores. (C) Spectra from different fragmentation techniques were also investigated. (D) For the ETD DMTm result, three different values each for fragment tolerance, fit score, and signal-to-noise cutoff were used to iterate through all combinations in order to find an optimized parameter set that balances the percent of decoy matches with the number of fragment ion matches observed for the true proteoform.

Interestingly, while the impact of settings on the number of expected decoy matches was substantial for all three parameters, there was relatively minimal impact on the number of total fragment matches with the proteoform. For example, increasing the S/N threshold from 3 to 20 led to a 2.5-fold decrease in decoy matches while only reducing the number of matched fragments by ∼5% (**Figure 4D**). Changing multiple parameters simultaneously, we were able to find a set of parameters that significantly cut the average number of decoy matches while retaining a high number of fragment ions (**Figure 4D**, optimized data points). The decoy matches were reduced by 4.1-fold to 3.9% and 2.2% expected decoy matches for shuffled and random sequences, with a less than 6% change in the total number of matching fragment ions. These parameter optimizations move toward automated analyses as significant reductions in decoy matches reduce the need for in-depth manual validation, except for validation of important sequence regions (e.g., key fragment ions supporting modification localization).

### Top-down sequencing of therapeutic antibodies

Antibody sequencing plays a central role not only in therapeutic development and quality control but also in clinical and biomedical research.^30^ We therefore sought to apply DMTm-enabled deep proteoform sequencing to antibodies. We chose SILuMab as a benchmark antibody to evaluate the top-down DMTm with ETD fragmentation that produced the most comprehensive sequencing of the standard proteins. We achieved 83.9% and 74.8% sequence coverage for the Lc and deglycosylated Hc, respectively (**Figure S12-13**). Following the addition of ensemble data to the DMTm results, we improved the sequence coverage to 87.6% and 77.5% for Lc and Hc. Here, ensemble antibody sequencing data had limited detection of larger fragment ions. In contrast, DMTm analysis of Hc produced 43 matching fragment ions with molecular weights above 30 kDa and seven matching fragment ions above 40 kDa, allowing confident fragment mapping for key regions in both subunits. For the Hc, 86.6% sequence coverage of the middle 350 residues of the chain (residues 50-399) was achieved. Without considering the N-terminal sides of the 28 proline residues, >90% regional sequence coverage was achieved. For the Lc, 98.3% of the inner residues that were at least 50 residues from each terminus (residues 50-168) were cleaved.

These results also increase fragment mapping confidence due to the identification of complementary, bidirectional ions. In the Lc data, 60.1% of matched ions were part of a complementary pair of *c*- and *z*-ions, with 131 total pairs formed. For the larger Hc subunit, 52 complementary pairs were formed. Although these complementary ions do not directly enhance sequence coverage, they help confirm fragment identities and improve overall confidence in the mapping. Accordingly, complementary ions could greatly assist in future de novo sequencing efforts.

### Automation and Increasing Throughput

As the data above relied on long acquisition times (e.g., 3 h for SILuMab Hc), we sought to achieve higher-throughput data collection while maintaining the same high-performance protein characterization. First, we moved the analysis to the newest generation instrument, the Thermo Scientific Orbitrap Tribrid Apex mass spectrometer. Secondly, we implemented an updated version of Advanced Ion Control (AIC) to further increase the number of ions per scan to maximize the amount of information obtainable per unit time while still maintaining individual ion levels.^31^ We collected an average of 4,381 ± 446 ions per spectrum with 3,054 ± 703 ions charge assigned per spectrum. The number of charge-assigned ions was ∼7-fold higher per spectrum than the SILuMab experiments above (**Figure S14**).

We achieved 98.1% and 83.3% manually validated sequence coverage for the Lc and Hc, respectively, from a 10-minute top-down DMTm acquisition with IR-EThcD activation applied to each subunit of disulfide-reduced NISTmAb (**Figure 5**). We observed an abundant number of bidirectional fragment pairs, with 157 for Lc and 142 for Hc. For residues greater than 50 amino acids away from each terminus (i.e., the inner residues of the protein), 100% and >95% were fragmented for the Lc and Hc, respectively. We also acquired fragmentation data in ensemble mode. After aggregation of the results in Proteoform Studio, 90.2% sequence coverage was achieved for the Hc (Lc sequence coverage was unchanged).

**Figure 5.**
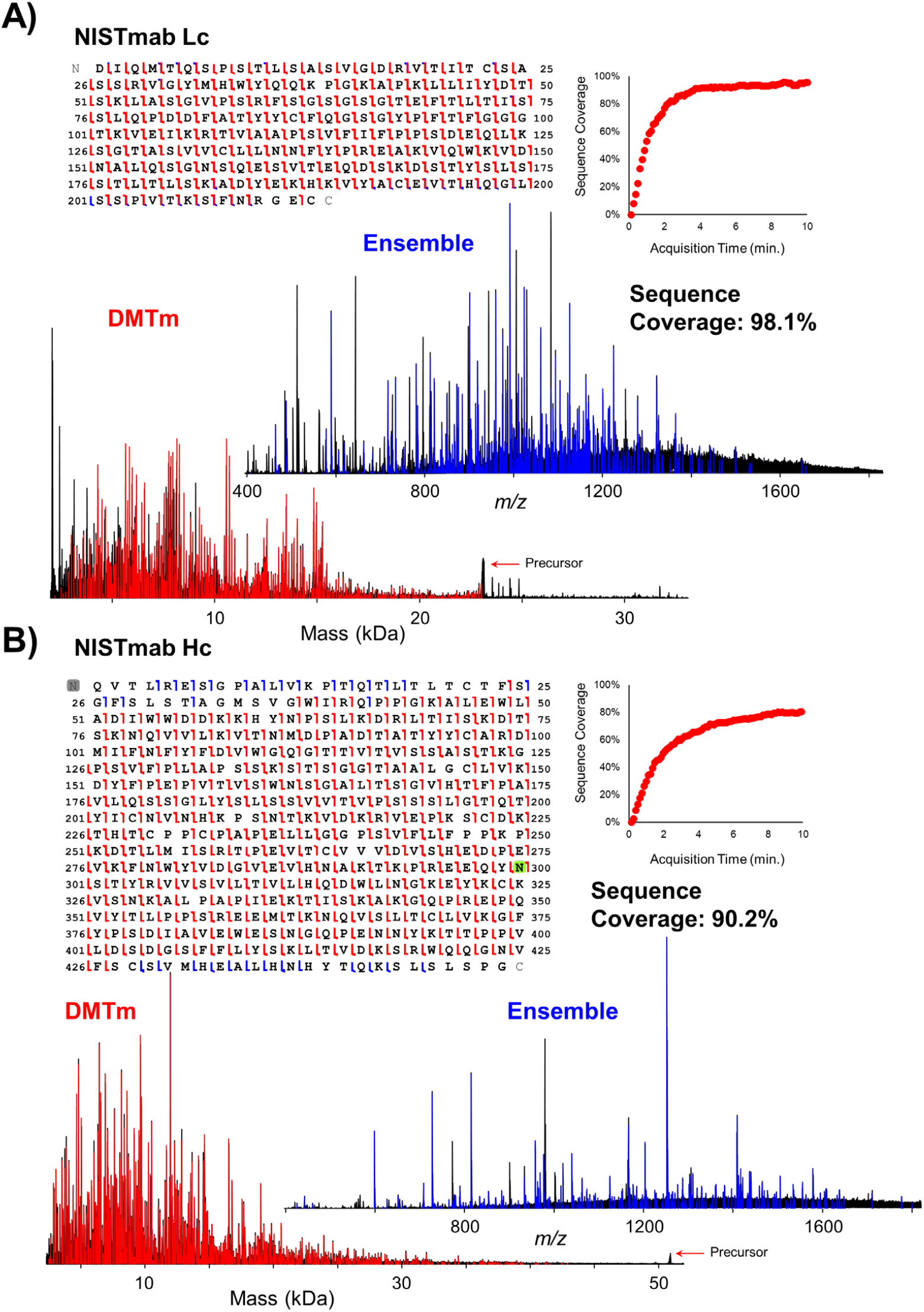
NISTmAb sequencing of Lc and Hc using IR-EThcD on a Thermo Scientific Orbitrap Tribrid Apex mass spectrometer. (A) Lc and (B) Hc were both analyzed with IR-EThcD for 10 minutes using both ensemble and DMTm. Data were combined with Proteoform Studio to show aggregated sequence coverage and fragmentation maps. Blue ions are from ensemble analysis, and red ions are from DMTm analysis.

We also further developed the Proteoform Studio platform to provide real-time feedback as the data collection progressed. By batching the number of ions that are sent to the voting algorithm, the rate of processing on a standard instrument computer is faster than the acquisition rate, enabling real-time metrics to be populated. We monitored the acquisition to assess when the sequence coverage had plateaued (**Figure 5, right side**). We also applied an auto-calibration procedure to automatically find the optimal calibration for the current spectrum. After applying the calibration shift to the spectra, the ppm tolerance of the isotopic fitter was tightened by using 3× median absolute deviation (MAD). For this NISTmAb data, Proteoform Studio automatically set window tolerances of 0.92 ppm for the Lc and 0.75 ppm for the Hc, producing decoy percentages for random sequences of 1.4% and 1.7% when matching against the four main ion types (*b, y, c*, and *z*) while only slightly decreasing sequence coverage from the manually validated levels (92.9% for Lc and 72.5% for Hc). Overall, these sequence coverage results paired with low false-discovery rates are step-function improvements over existing top-down capabilities and highlight the potential for deep proteoform sequencing to be performed in a high-throughput, automated fashion using this complete workflow.

## Conclusion

Our study provides the first comprehensive workflow for deep proteoform sequencing using top-down DMTm with the different fragmentation techniques available on modern Orbitrap Tribrid systems. With the automated data processing of Proteoform Studio, the entire process is significantly simplified and could be readily adopted in any lab with such instrumentation. We evaluated top-down DMTm performance for intact protein characterization across HCD, ETD, UVPD, and IR-EThcD fragmentation methods, establishing a benchmark for future applications of the technique. Through direct mass measurement of individual ions, top-down DMTm effectively circumvents much of the spectral congestion historically observed at the *m/z* spectrum level, enabling detection of more fragment ions, different types of fragment ions, and higher-mass fragment ions that are rarely accessible with conventional top-down measurements. By integrating DMTm results with ensemble results in Proteoform Studio, we were able to combine the strengths of both approaches, achieving unprecedentedly high sequence coverage of disulfide-reduced antibodies.

Such high accuracy and sensitivity positions top-down DMTm as a powerful platform for next-generation top-down proteomics, with strong potential for the development of *de novo* sequencing, detection of single amino acid variants, and precise localization of multiple PTMs on intact proteins.^39^ The deep sequencing capabilities with top-down DMTm for larger proteins advance proteoform characterization and opens up new applications that previously would have had a low probability of success. Many biologically and pharmaceutically relevant proteins are larger than 30 kDa and have historically not been successfully characterized using routine top-down analysis methods. With the extensive sequence coverage offered by DMTm, particularly in the middle regions of larger proteins, studying these large proteins will become more tractable.

Coupled with the extremely low sample requirements for DMTm, a much larger dynamic range of proteoforms can be accessed and sequenced. By unlocking high mass fragment ions, future workflows could utilize top-down for confirmation of chain pairing with multispecifics and even enable fully intact antibody analyses without the need for any digestion. Through continued development of instrumentation and data analysis pipelines, dissemination of the technology, and improved acquisition throughput, top-down DMTm will play a central role in driving high-resolution proteoform characterization, advancing the applications of top-down mass spectrometry in biopharmaceutical analyses and clinical biomarker discovery.

## Acknowledgements

This work was supported by the NIGMS of the National Institutes of Health under award numbers R44 GM136046 and RM1 GM156535. The content is solely the responsibility of the authors and does not necessarily represent the official views of the National Institutes of Health. The authors thank Mike Senko, Ping Yip, Kyle Bowen, Mike Goodwin, Rafa Melani, Joe Greer, Lily Miller, and Matt Robey for helpful discussions and contributions to the development of top-down Direct Mass Technology and the Proteoform Studio platform.

## Notes

### Competing Interest Statement

K.R.D. and R.T.F. are employed by Thermo Fisher Scientific, a company that commercializes mass spectrometers and bioinformatics software.

## References

(1) Smith, L. M.; Kelleher, N. L.; Proteomics, C. f. T. D. Proteoform: a single term describing protein complexity. Nat Methods 2013, 10 (3), 186–187. DOI: 10.1038/nmeth.2369.

(2) Aebersold, R.; Agar, J. N.; Amster, I. J.; Baker, M. S.; Bertozzi, C. R.; Boja, E. S.; Costello, C. E.; Cravatt, B. F.; Fenselau, C.; Garcia, B. A.; et al. How many human proteoforms are there? Nat Chem Biol 2018, 14 (3), 206–214. DOI: 10.1038/nchembio.2576.

(3) Roberts, D. S.; Loo, J. A.; Tsybin, Y. O.; Liu, X.; Wu, S.; Chamot-Rooke, J.; Agar, J. N.; Paša-Tolić, L.; Smith, L. M.; Ge, Y. Top-down proteomics. Nat Rev Methods Primers 2024, 4 (1). DOI: 10.1038/s43586-024-00318-2.

(4) Millán-Martín, S.; Jakes, C.; Carillo, S.; Gallagher, L.; Scheffler, K.; Broster, K.; Bones, J. Multi-Attribute Method (MAM): An Emerging Analytical Workflow for Biopharmaceutical Characterization, Batch Release and cGMP Purity Testing at the Peptide and Intact Protein Level. Crit Rev Anal Chem 2024, 54 (8), 3234–3251. DOI: 10.1080/10408347.2023.2238058.

(5) Po, A.; Eyers, C. E. Top-Down Proteomics and the Challenges of True Proteoform Characterization. J Proteome Res 2023, 22 (12), 3663–3675. DOI: 10.1021/acs.jproteome.3c00416.

(6) Zhou, M.; Lantz, C.; Brown, K. A.; Ge, Y.; Paša-Tolić, L.; Loo, J. A.; Lermyte, F. Higher-order structural characterisation of native proteins and complexes by top-down mass spectrometry. Chem Sci 2020, 11 (48), 12918–12936. DOI: 10.1039/d0sc04392c.

(7) Aebersold, R.; Mann, M. Mass spectrometry-based proteomics. Nature 2003, 422 (6928), 198–207. DOI: 10.1038/nature01511.

(8) Catherman, A. D.; Skinner, O. S.; Kelleher, N. L. Top Down proteomics: facts and perspectives. Biochem Biophys Res Commun 2014, 445 (4), 683–693. DOI: 10.1016/j.bbrc.2014.02.041.

(9) Durbin, K. R.; Fornelli, L.; Fellers, R. T.; Doubleday, P. F.; Narita, M.; Kelleher, N. L. Quantitation and Identification of Thousands of Human Proteoforms below 30 kDa. J Proteome Res 2016, 15 (3), 976–982. DOI: 10.1021/acs.jproteome.5b00997.

(10) Brodbelt, J. S. Deciphering combinatorial post-translational modifications by top-down mass spectrometry. Curr Opin Chem Biol 2022, 70, 102180. DOI: 10.1016/j.cbpa.2022.102180.

(11) Donnelly, D. P.; Rawlins, C. M.; DeHart, C. J.; Fornelli, L.; Schachner, L. F.; Lin, Z.; Lippens, J. L.; Aluri, K. C.; Sarin, R.; Chen, B.; et al. Best practices and benchmarks for intact protein analysis for top-down mass spectrometry. Nat Methods 2019, 16 (7), 587–594. DOI: 10.1038/s41592-019-0457-0.

(12) Tabb, D. L.; Jeong, K.; Druart, K.; Gant, M. S.; Brown, K. A.; Nicora, C.; Zhou, M.; Couvillion, S.; Nakayasu, E.; Williams, J. E.; et al. Comparing Top-Down Proteoform Identification: Deconvolution, PrSM Overlap, and PTM Detection. J Proteome Res 2023, 22 (7), 2199–2217. DOI: 10.1021/acs.jproteome.2c00673.

(13) Khristenko, N. A.; Nagornov, K. O.; Garcia, C.; Gasilova, N.; Gant, M.; Druart, K.; Kozhinov, A. N.; Menin, L.; Chamot-Rooke, J.; Tsybin, Y. O. Top-Down and Middle-Down Mass Spectrometry of Antibodies. Mol Cell Proteomics 2025, 24 (7), 100989. DOI: 10.1016/j.mcpro.2025.100989.

(14) Kline, J. T.; Melani, R. D.; Fornelli, L. Mass spectrometry characterization of antibodies at the intact and subunit levels: from targeted to large-scale analysis. Int J Mass Spectrom 2023, 492. DOI: 10.1016/j.ijms.2023.117117.

(15) Drown, B. S.; Gupta, R.; McGee, J. P.; Hollas, M. A. R.; Hergenrother, P. J.; Kafader, J. O.; Kelleher, N. L. Precise Readout of MEK1 Proteoforms upon MAPK Pathway Modulation by Individual Ion Mass Spectrometry. Anal Chem 2024, 96 (11), 4455–4462. DOI: 10.1021/acs.analchem.3c04758.

(16) Olsen, J. V.; Macek, B.; Lange, O.; Makarov, A.; Horning, S.; Mann, M. Higher-energy C-trap dissociation for peptide modification analysis. Nat Methods 2007, 4 (9), 709–712. DOI: 10.1038/nmeth1060.

(17) Kim, M. S.; Pandey, A. Electron transfer dissociation mass spectrometry in proteomics. Proteomics 2012, 12 (4-5), 530–542. DOI: 10.1002/pmic.201100517.

(18) Brodbelt, J. S.; Morrison, L. J.; Santos, I. Ultraviolet Photodissociation Mass Spectrometry for Analysis of Biological Molecules. Chem Rev 2020, 120 (7), 3328–3380. DOI: 10.1021/acs.chemrev.9b00440.

(19) Durbin, K. R.; Skinner, O. S.; Fellers, R. T.; Kelleher, N. L. Analyzing internal fragmentation of electrosprayed ubiquitin ions during beam-type collisional dissociation. J Am Soc Mass Spectrom 2015, 26 (5), 782–787. DOI: 10.1007/s13361-015-1078-1.

(20) Mikawy, N. N.; Rojas Ramírez, C.; DeFiglia, S. A.; Szot, C. W.; Le, J.; Lantz, C.; Wei, B.; Zenaidee, M. A.; Blakney, G. T.; Nesvizhskii, A. I.; et al. Are Internal Fragments Observable in Electron Based Top-Down Mass Spectrometry? Mol Cell Proteomics 2024, 23 (9), 100814. DOI: 10.1016/j.mcpro.2024.100814.

(21) Madsen, J. A.; Boutz, D. R.; Brodbelt, J. S. Ultrafast ultraviolet photodissociation at 193 nm and its applicability to proteomic workflows. J Proteome Res 2010, 9 (8), 4205–4214. DOI: 10.1021/pr100515x.

(22) Kessler, A. L.; Fort, K. L.; Resemann, H. C.; Krüger, P.; Wang, C.; Koch, H.; Hauschild, J. P.; Marino, F.; Heck, A. J. R. Increased EThcD Efficiency on the Hybrid Orbitrap Excedion Pro Mass Analyzer Extends the Depth in Identification and Sequence Coverage of HLA Class I Immunopeptidomes. Mol Cell Proteomics 2025, 24 (9), 101049. DOI: 10.1016/j.mcpro.2025.101049.

(23) Carfagno, A. K.; Lieu, L. B.; Kline, J. T.; Fornelli, L.; Durbin, K. R. Automating Middle-Down Mass Spectrometry Analysis for Extensive Antibody Characterization. Anal Chem 2026, 98 (16), 11707–11719. DOI: 10.1021/acs.analchem.5c06408.

(24) Dunham, S. D.; Juetten, K. J.; Hellinger, J.; Gadallah, M. I.; Dioli, O. E.; Brodbelt, J. S. Leveraging Complementary Ion Activation Methods with Proton Transfer Charge Reduction Reactions for Comprehensive Characterization of Monoclonal Antibody Heavy Chain Subunits. Anal Chem 2025, 97 (27), 14281–14289. DOI: 10.1021/acs.analchem.5c01075.

(25) Kafader, J. O.; Melani, R. D.; Durbin, K. R.; Ikwuagwu, B.; Early, B. P.; Fellers, R. T.; Beu, S. C.; Zabrouskov, V.; Makarov, A. A.; Maze, J. T.; et al. Multiplexed mass spectrometry of individual ions improves measurement of proteoforms and their complexes. Nat Methods 2020, 17 (4), 391–394. DOI: 10.1038/s41592-020-0764-5.

(26) Kafader, J. O.; Beu, S. C.; Early, B. P.; Melani, R. D.; Durbin, K. R.; Zabrouskov, V.; Makarov, A. A.; Maze, J. T.; Shinholt, D. L.; Yip, P. F.; et al. STORI Plots Enable Accurate Tracking of Individual Ion Signals. J Am Soc Mass Spectrom 2019, 30 (11), 2200–2203. DOI: 10.1007/s13361-019-02309-0.

(27) Du, C.; Cotham, V. C.; Thakur, G.; Mink, S.; Tustian, A. D.; Møller-Tank, S.; Wang, S.; Li, N. An Online Buffer Exchange Platform for Charge Detection Mass Spectrometry Analysis of AAVs and AAV-Antibody Complexes. J Am Soc Mass Spectrom 2025, 36 (10), 2230–2238. DOI: 10.1021/jasms.5c00206.

(28) Kafader, J. O.; Durbin, K. R.; Melani, R. D.; Des Soye, B. J.; Schachner, L. F.; Senko, M. W.; Compton, P. D.; Kelleher, N. L. Individual Ion Mass Spectrometry Enhances the Sensitivity and Sequence Coverage of Top-Down Mass Spectrometry. J Proteome Res 2020, 19 (3), 1346–1350. DOI: 10.1021/acs.jproteome.9b00797.

(29) Xu, T.; Su, T.; Soye, B. J. D.; Kandi, S.; Huang, C. F.; Wilkins, J. T.; Castellani, R. J.; Kafader, J. O.; Patrie, S. M.; Vassar, R.; et al. The Proteoform Landscape of Tau from the Human Brain. J Proteome Res 2025, 24 (6), 2916–2925. DOI: 10.1021/acs.jproteome.5c00139.

(30) de Graaf, S. C.; Hoek, M.; Tamara, S.; Heck, A. J. R. A perspective toward mass spectrometry-based de novo sequencing of endogenous antibodies. MAbs 2022, 14 (1), 2079449. DOI: 10.1080/19420862.2022.2079449.

(31) McGee, J. P.; Senko, M. W.; Jooß, K.; Des Soye, B. J.; Compton, P. D.; Kelleher, N. L.; Kafader, J. O. Automated Control of Injection Times for Unattended Acquisition of Multiplexed Individual Ion Mass Spectra. Anal Chem 2022, 94 (48), 16543–16548. DOI: 10.1021/acs.analchem.2c03495.

