## Supplemental Info for "Deep Proteoform Sequencing with Top-Down Direct Mass Technology"

### Composite data contain limited unique fragment information

Composite data is defined here as the time-averaged *m/z* spectrum acquired during DMTm data acquisition. Since it is an *m/z* spectrum like the spectra from conventional ensemble acquisition, a key question is whether such composite data can be processed similarly to ensemble data to extract unique sequence information, particularly lowly charged fragment ions near the termini. To investigate, we compared $MS^2$ spectra obtained from composite, ensemble, and DMTm acquisition of carbonic anhydrase (29 kDa). To induce production of additional lower mass fragment ions, the ETD reaction time was extended to 9 ms for ensemble mode, while composite data shared the same 1 ms ETD reaction time used in DMTm. As shown in **Figure S10A**, ensemble data yielded extensive terminal ions that almost completely covered the first and the last 18 amino acids of carbonic anhydrase. However, only four terminal ions were detected in these terminal regions from composite data. Insets in **Figure S10A** highlight terminal ions that were observed in the ensemble data but were either missing or had extremely weak signals in the composite data. The ions with <+4 charges are annotated in the ensemble *m/z* space because they cannot be detected in DMTm.[30] These results indicate that the ETD conditions optimized for DMTm acquisition do not generate sufficient terminal fragmentations. Thus, composite data does not substantially compensate for the low-charge ion detection limitation in DMTm. On the other hand, ensemble measurement under stronger activation conditions can easily and quickly generate many more terminal ions, suggesting additional ensemble acquisitions could be utilized as a complementary acquisition mode when extensive terminal coverage is not obtained by DMTm. This comparison also demonstrates that mild fragmentation conditions are more optimal for DMTm acquisition, as these conditions generate medium to large fragment ions that align well with DMTm detection.

Beyond terminal fragment ions, composite data also fails to provide additional sequence information of medium to large fragment ions compared to its corresponding DMTm data. **Figure S10B** illustrates a composite window ranging from 980 to 983 *m/z* that contains at least four fragment ions, as shown in the insets. Although these four ions were clearly observed in the DMTm spectra, their corresponding isotopic distributions were not discernible in the composite *m/z* domain. Furthermore, no charge variants of these ions were detectable across the entire range of composite *m/z* domain, indicating they were uniquely detected in the DMTm data type. Large fragment ions such as the ones in the inset produce wide isotopic distributions, often overlapping with one another and making composite spectrum uninterpretable. Even small fragments such as z51, while having less complex isotopic distributions, can also be masked by signals of other ions, especially if they are present at low abundance.

Overall, composite data provide mostly redundant sequence information already captured in the DMTm data. While composite spectra may reveal a small number of low-charge-state ions not captured by DMTm, these additional data are not significant enough to justify its use over ensemble measurements optimized to produce smaller terminal fragment ions. In addition, composite data interpretation requires extensive manual fragment annotation validation because the abundant precursor and ETnoD ions dominate the *m/z* spectrum and elevate the S/N threshold, further complicating automated fragment annotation in current software tools.[32,33] For practical purposes, collecting additional ensemble data optimized for obtaining terminal fragment ions is more feasible and effective than mining composite data for marginal gains on sequence coverage.

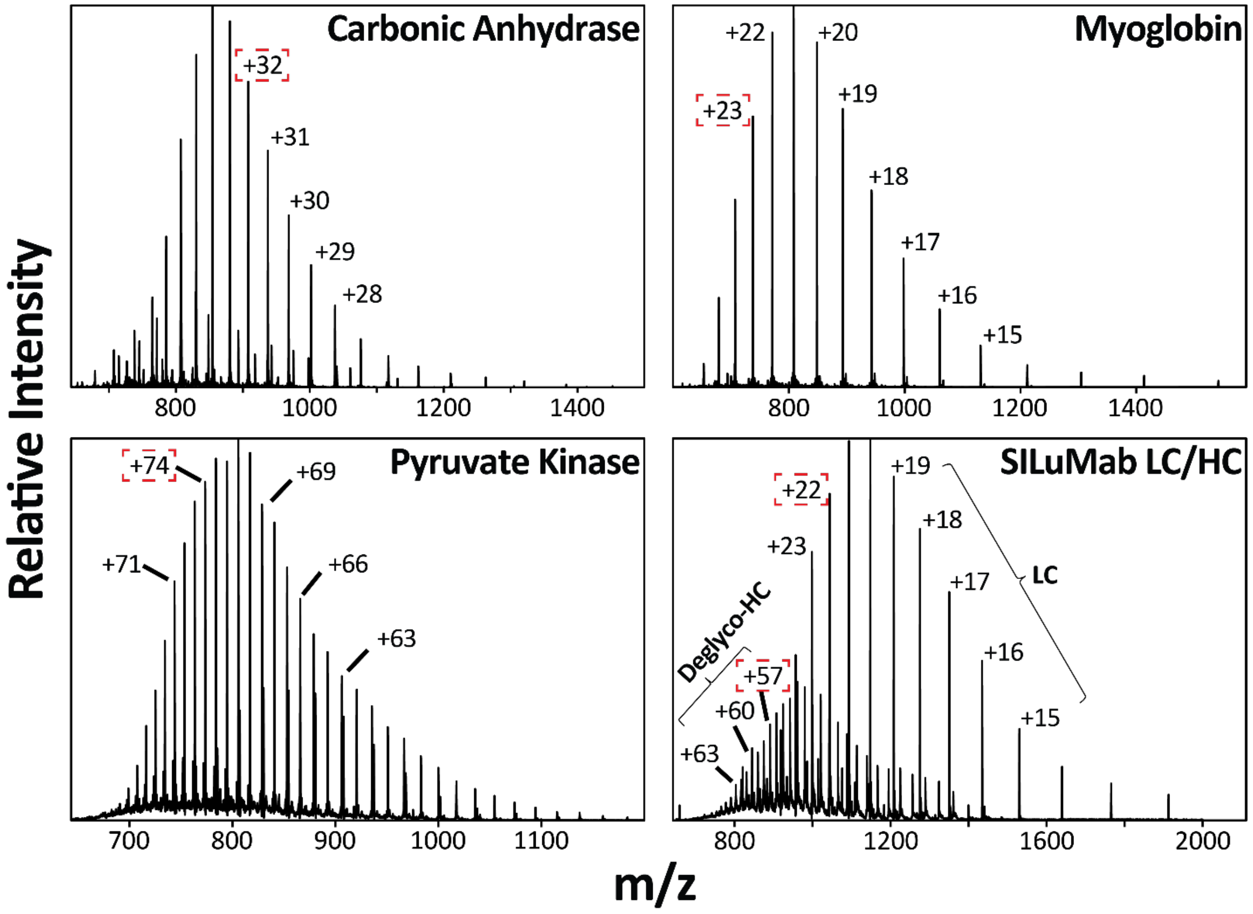


**Figure S1.** Charge distributions of three standard proteins and reduced and deglycosylated SILuMab. Carbonic anhydrase, myoglobin, pyruvate kinase, and SILuMab Lc and Hc subunits were subjected to intact mass profiling via ensemble measurement. Charge states highlighted by dashed red rectangles were selected for fragmentation.

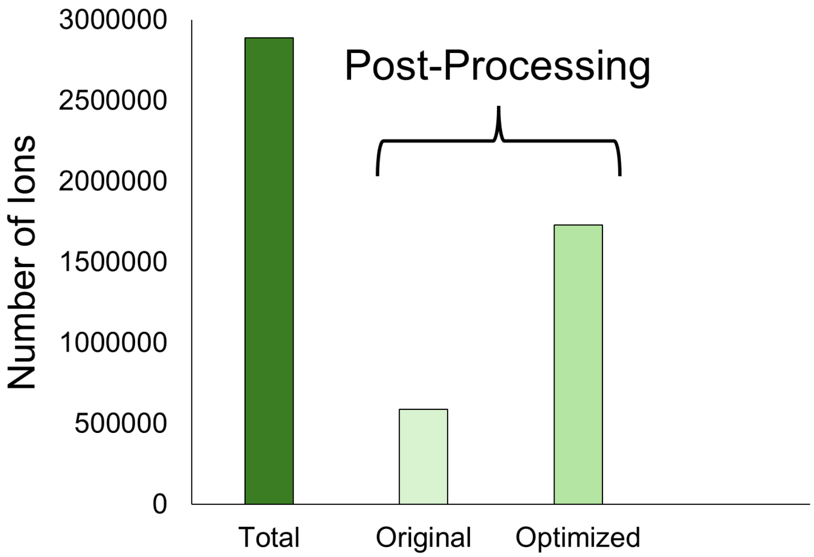


**Figure S2.** Ion counts for Direct Mass Technology mode (DMTm) processing. The total number of ions collected, the ions charge assigned using the original processing methodology, and the ions charge assigned using the optimized processing methodology developed here for top-down DMTm analyses are shown.

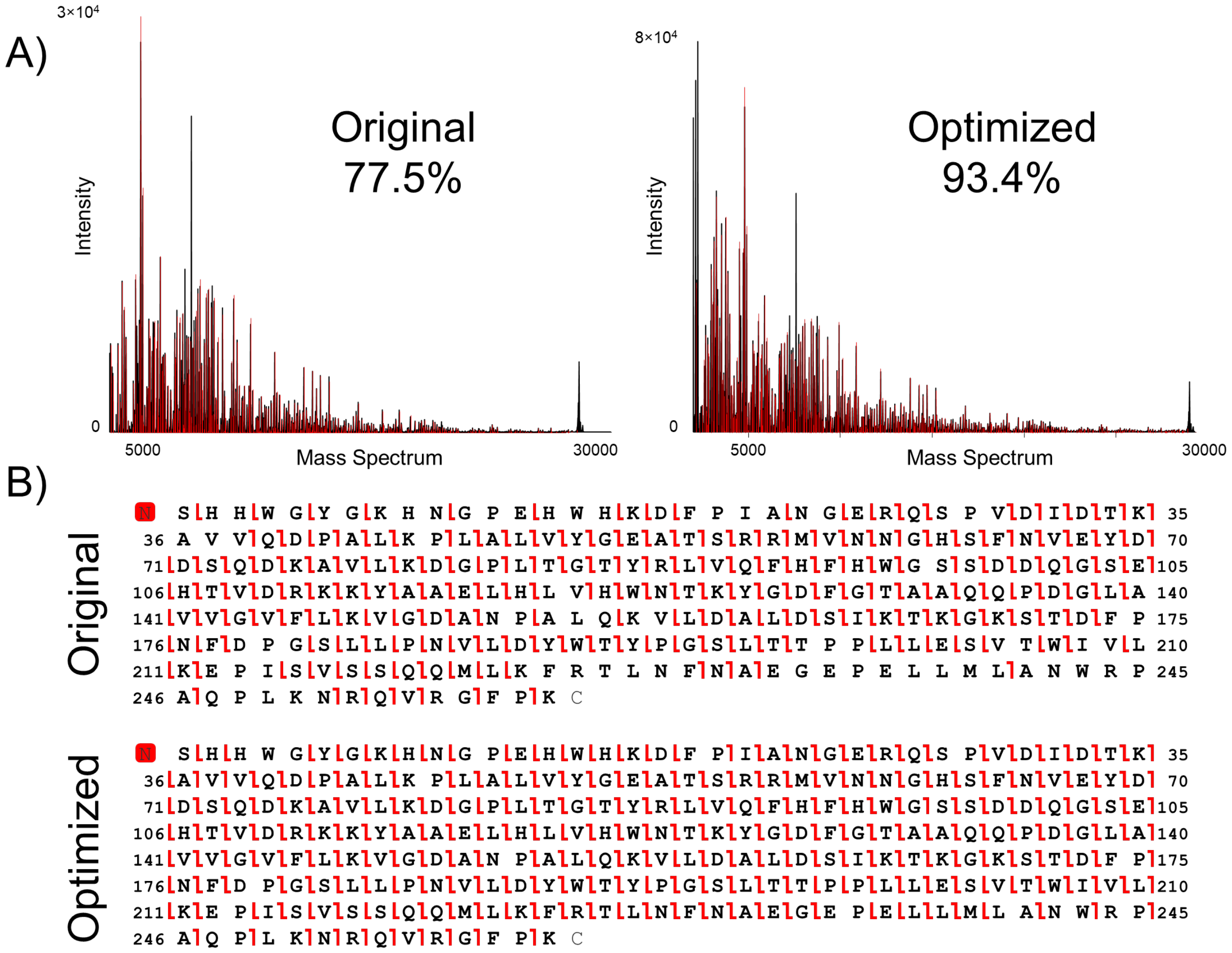


**Figure S3. Fragment ion coverages with two different DMTm processing and charge assignment routines.** (A) Spectra are shown with annotated distributions corresponding to matching fragment ions before (left) and after (right) optimization of DMTm individual ion processing. (B) The graphical fragment maps are also shown before (top) and (after) optimization of data processing routines.

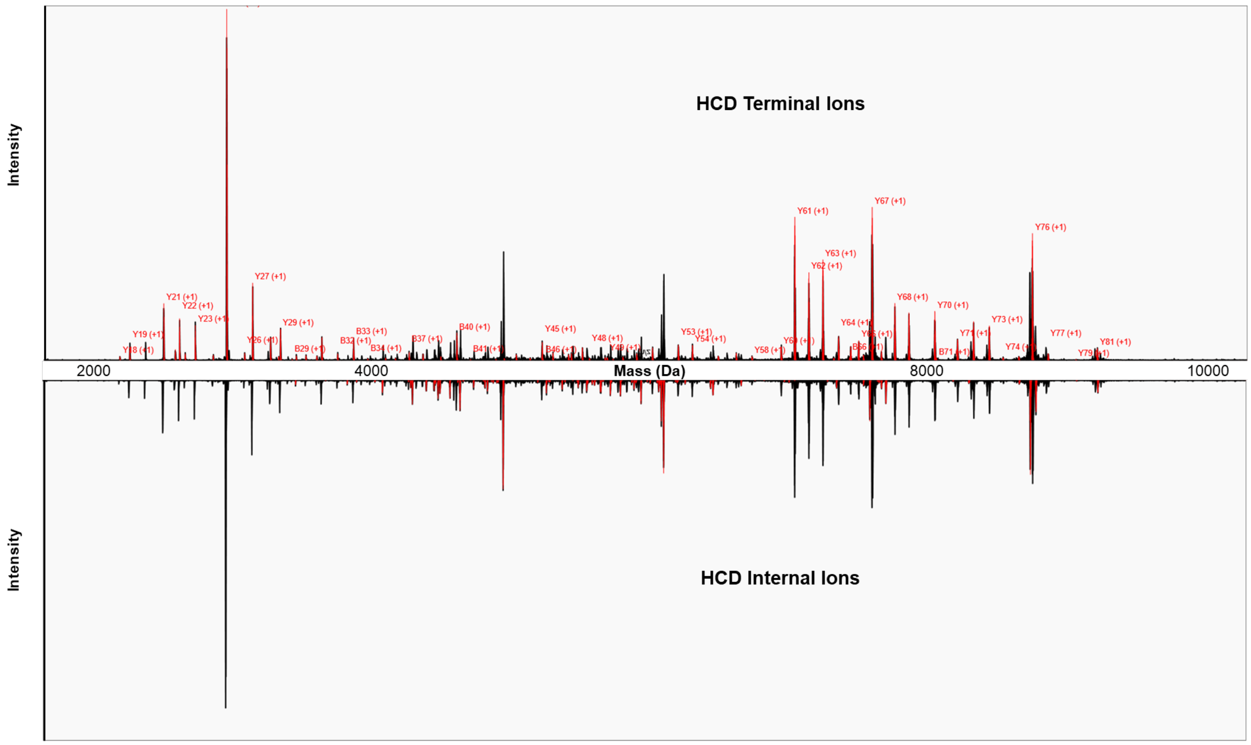


**Figure S4. Terminal and internal ions created by HCD fragmentation of carbonic anhydrase.** (Top) Matching terminal ions are annotated. (Bottom) Matching internal ions (*i.e.*, created by multiple fragmentation events) are displayed in red.

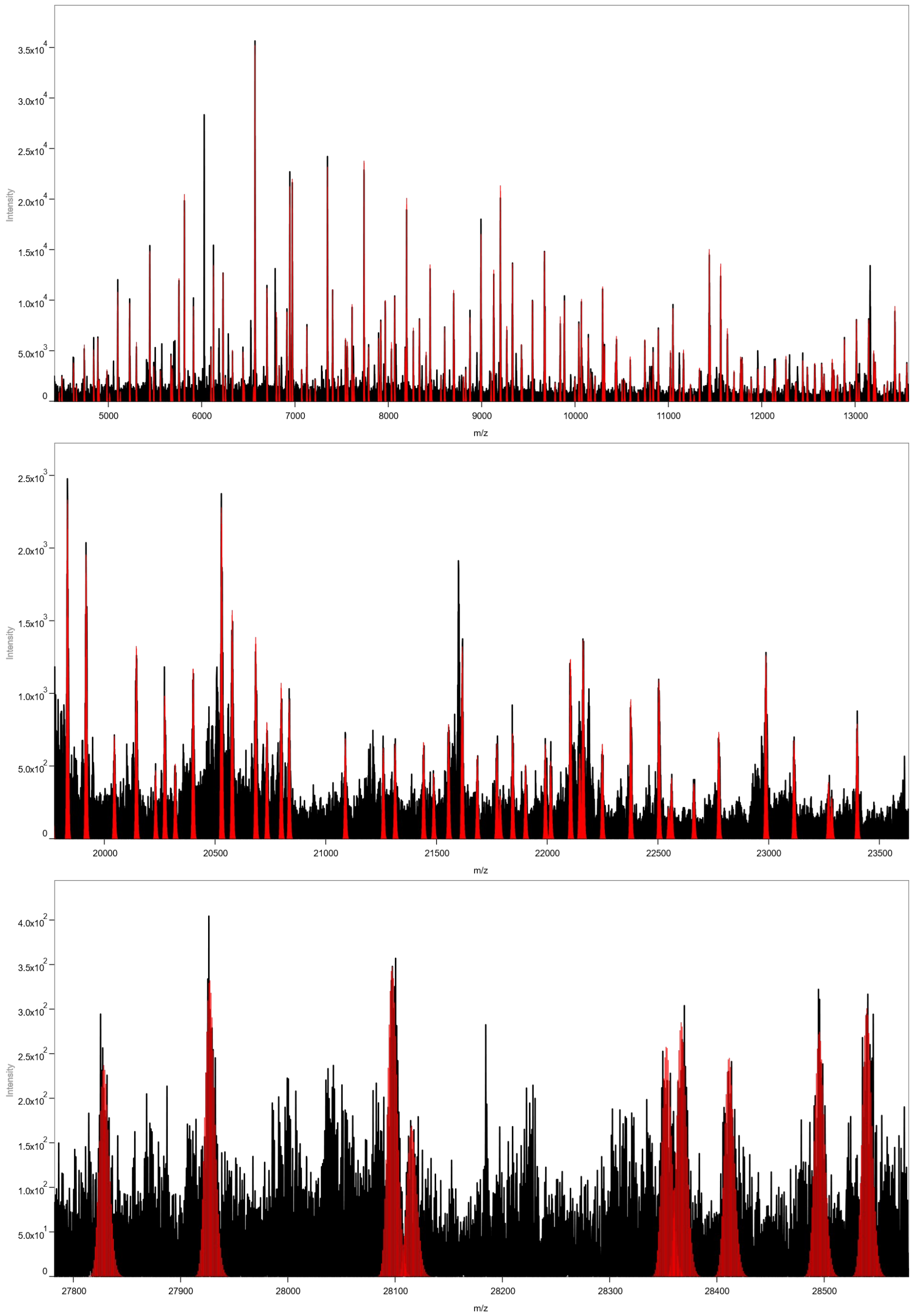


**Figure S5. DMTm charge assigned spectrum for pyruvate kinase with ETD.** Singly charged mass species (M+H) are shown for (A) 5000-13000 Da, (B) 20000-23500 Da, and (C) 27800-28600 Da.

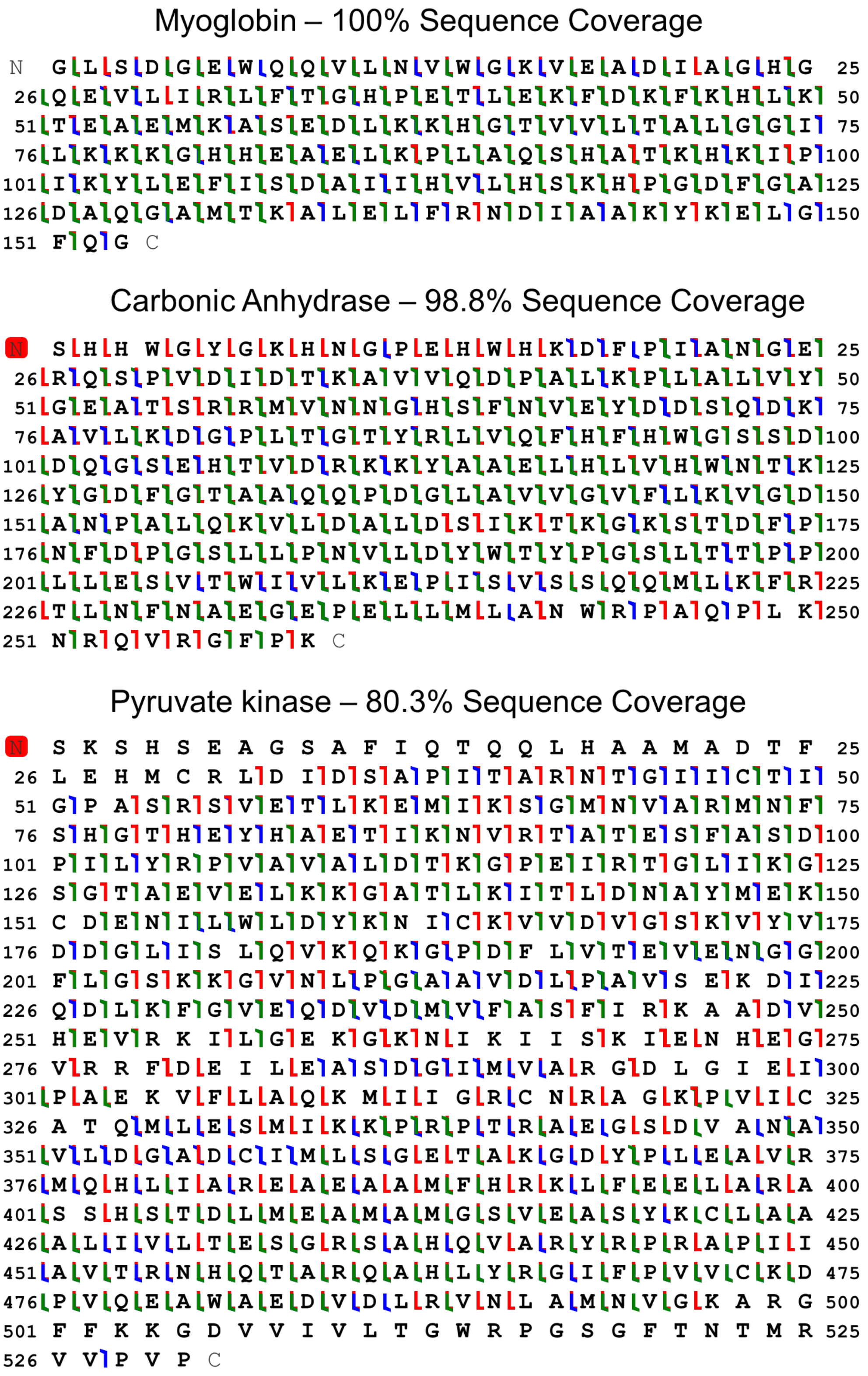


**Figure S6. Aggregated top-down DMTm fragmentation maps.** Matching fragment ions from DMTm ETD, HCD, and UVPD were aggregated together to form comprehensive fragment maps for (A) myoglobin, (B) carbonic anhydrase, and (C) pyruvate kinase. Blue fragment flags correspond to *b- and y*-ions, red fragment flags correspond to *c- and z*-ions, and green fragment flags correspond to *a*- and *x*-ions.

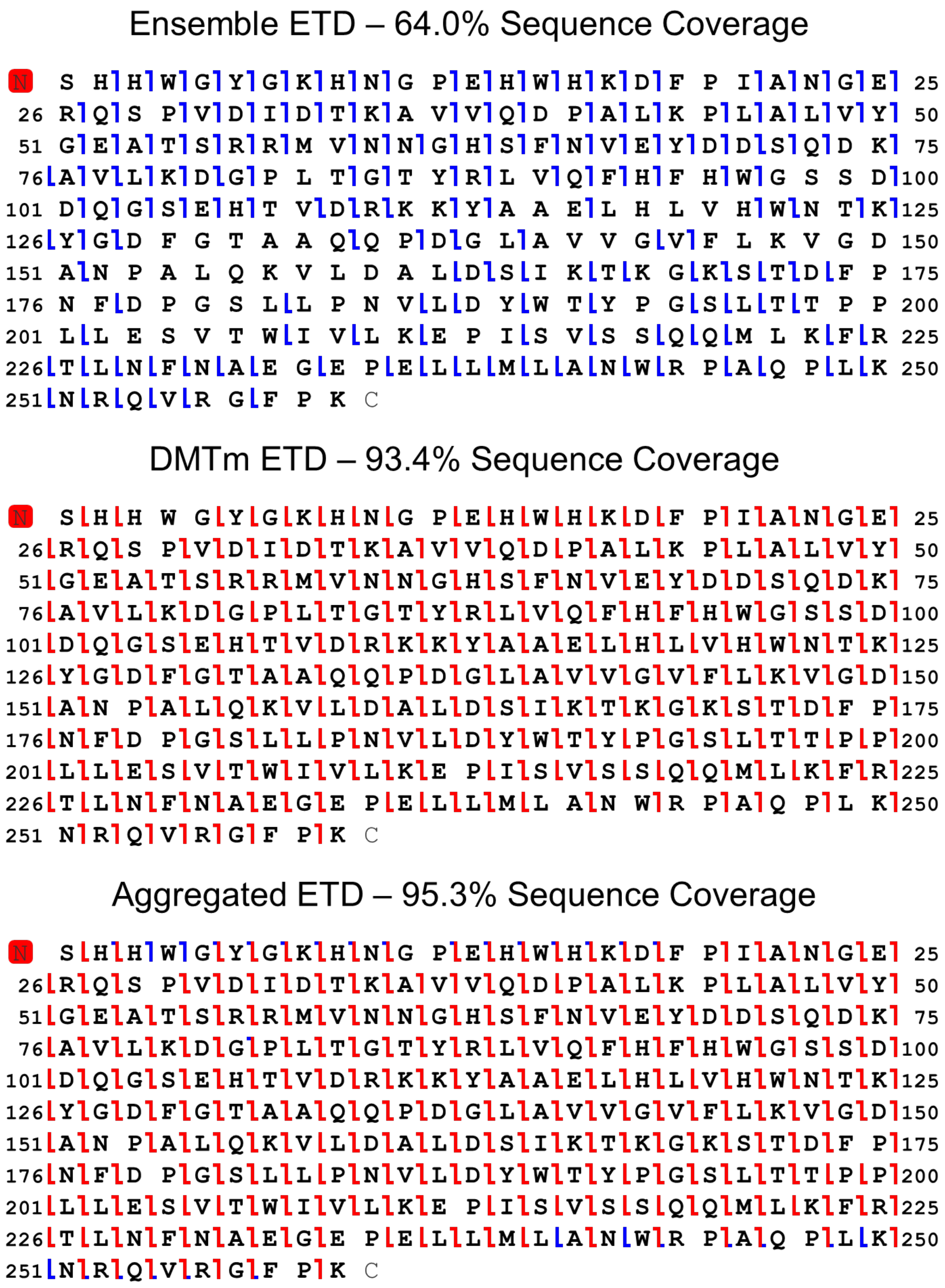


**Figure S7. Ensemble (top), DMTm (middle) and combined (bottom) ETD fragmentation maps for carbonic anhydrase.**

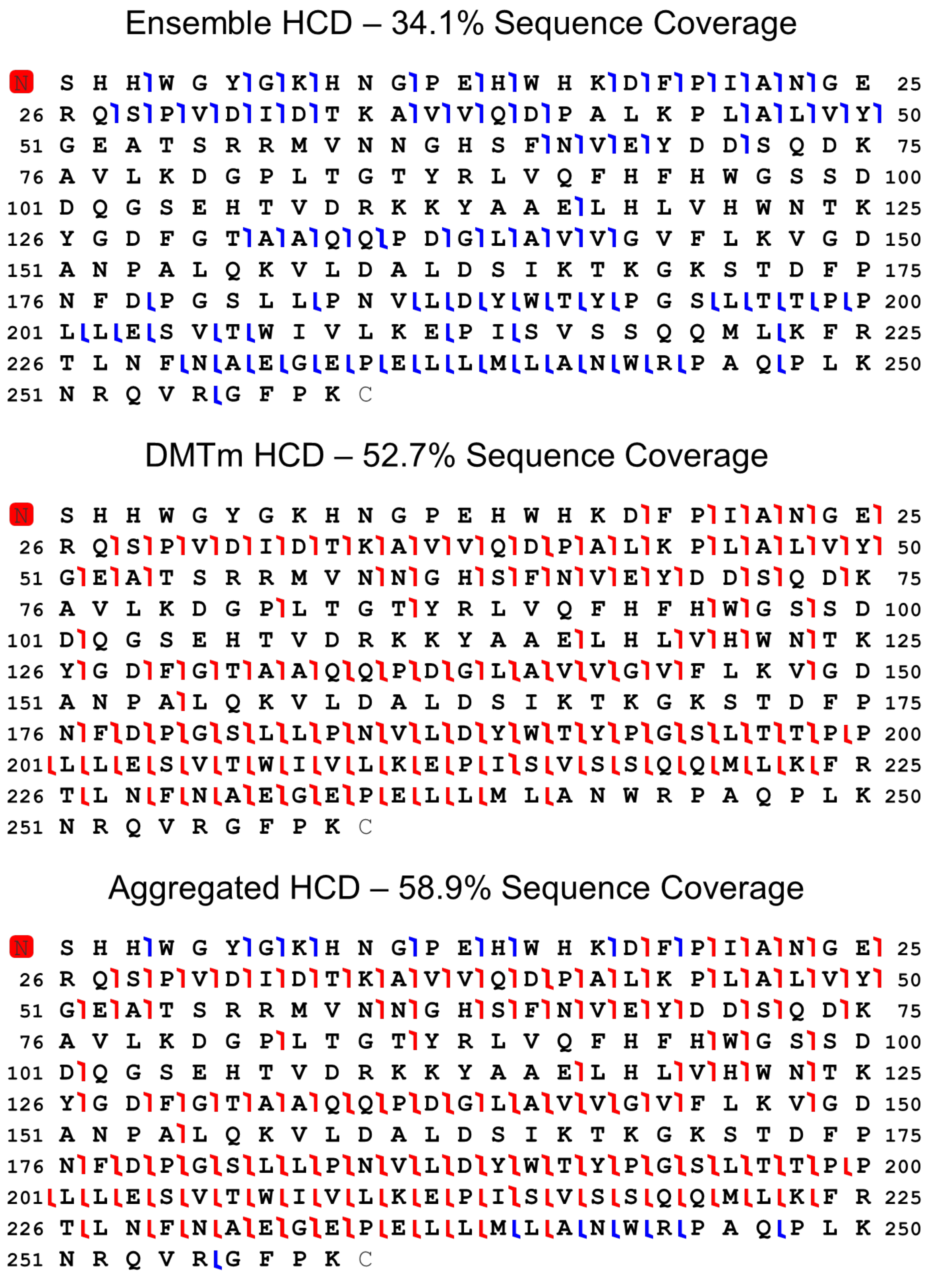


**Figure S8. Ensemble (top), DMTm (middle), and combined (bottom) HCD fragmentation maps for carbonic anhydrase.**

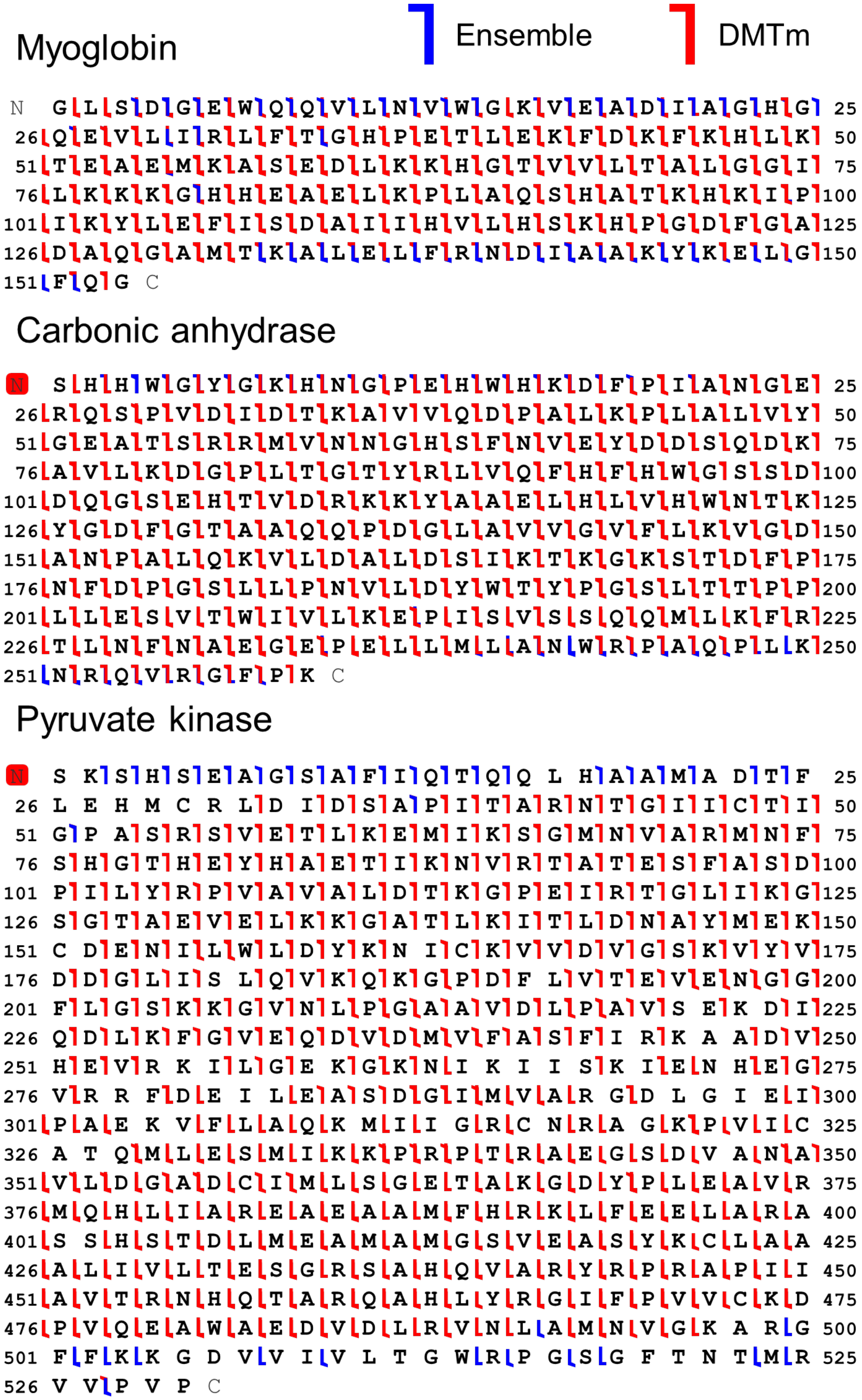


**Figure S9. Combined ensemble and DMTm graphical fragmentation maps for (top) myoglobin, (middle) carbonic anhydrase, and (bottom) pyruvate kinase.**

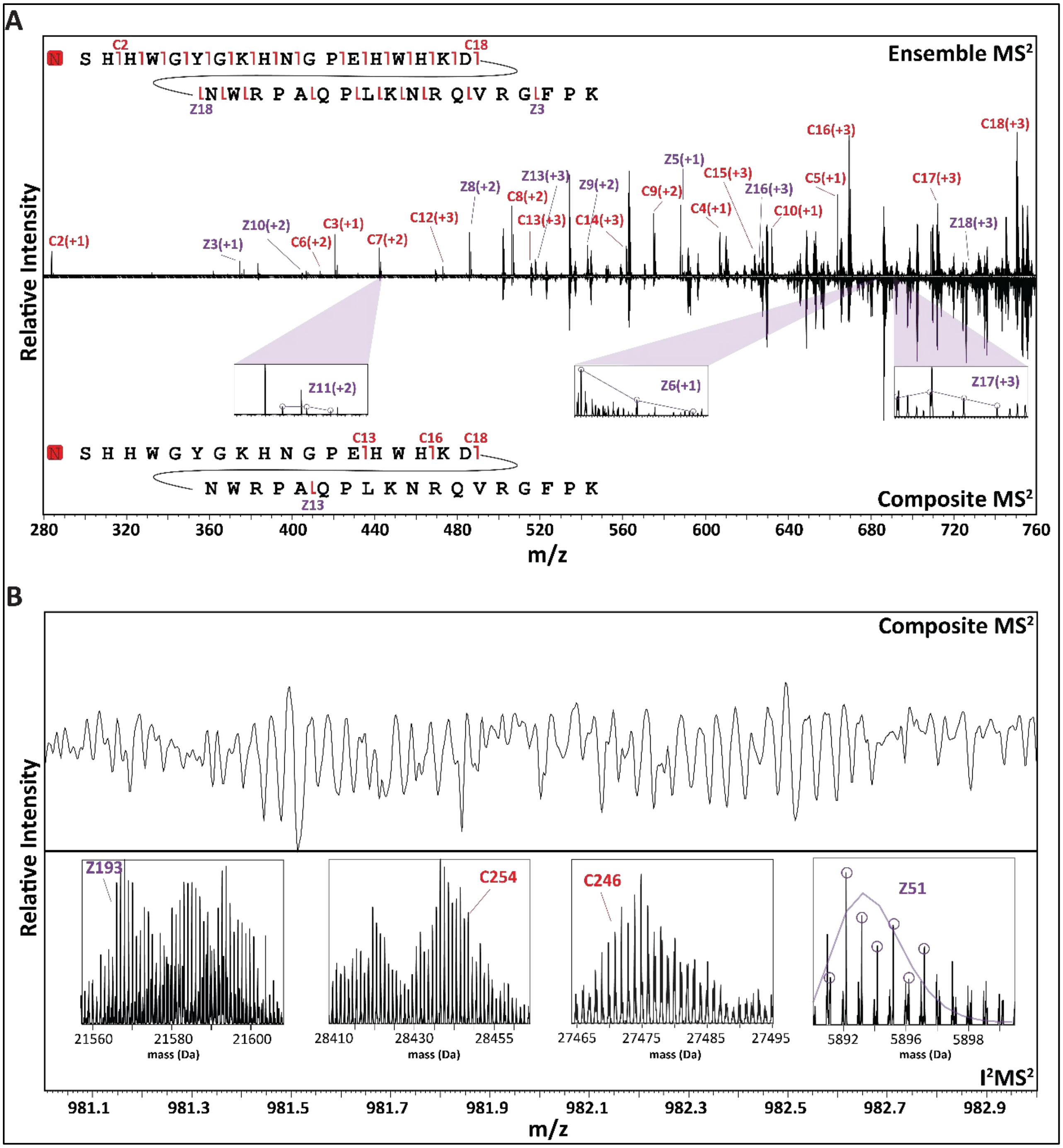


**Figure S10. Composite data contains less terminal ion information than ensemble data. (A)** Butterfly plot showing fragment ion annotations in the terminal regions of carbonic anhydrase for ensemble versus composite data. **(B)** Four fragment ions uniquely detected in DMTm data. Red ions indicate *c*-ions and purple ions indicate *z*-ions.

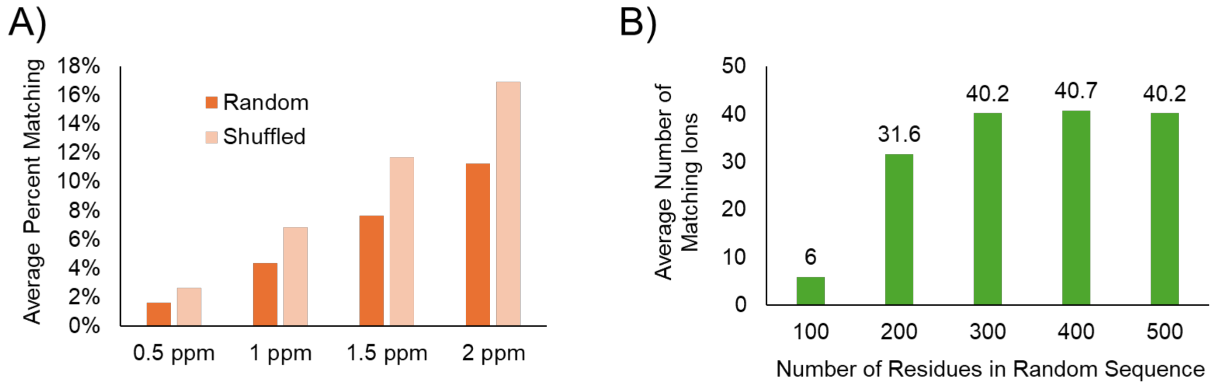


**Figure S11. Decoy matching fragment ions. (A)** The effect of changing the part-per-million (ppm) tolerance window for the isotopic fitter is shown for both randomized and shuffled sequences. **(B)** The impact of the number of residues on the number of matching ions is shown.

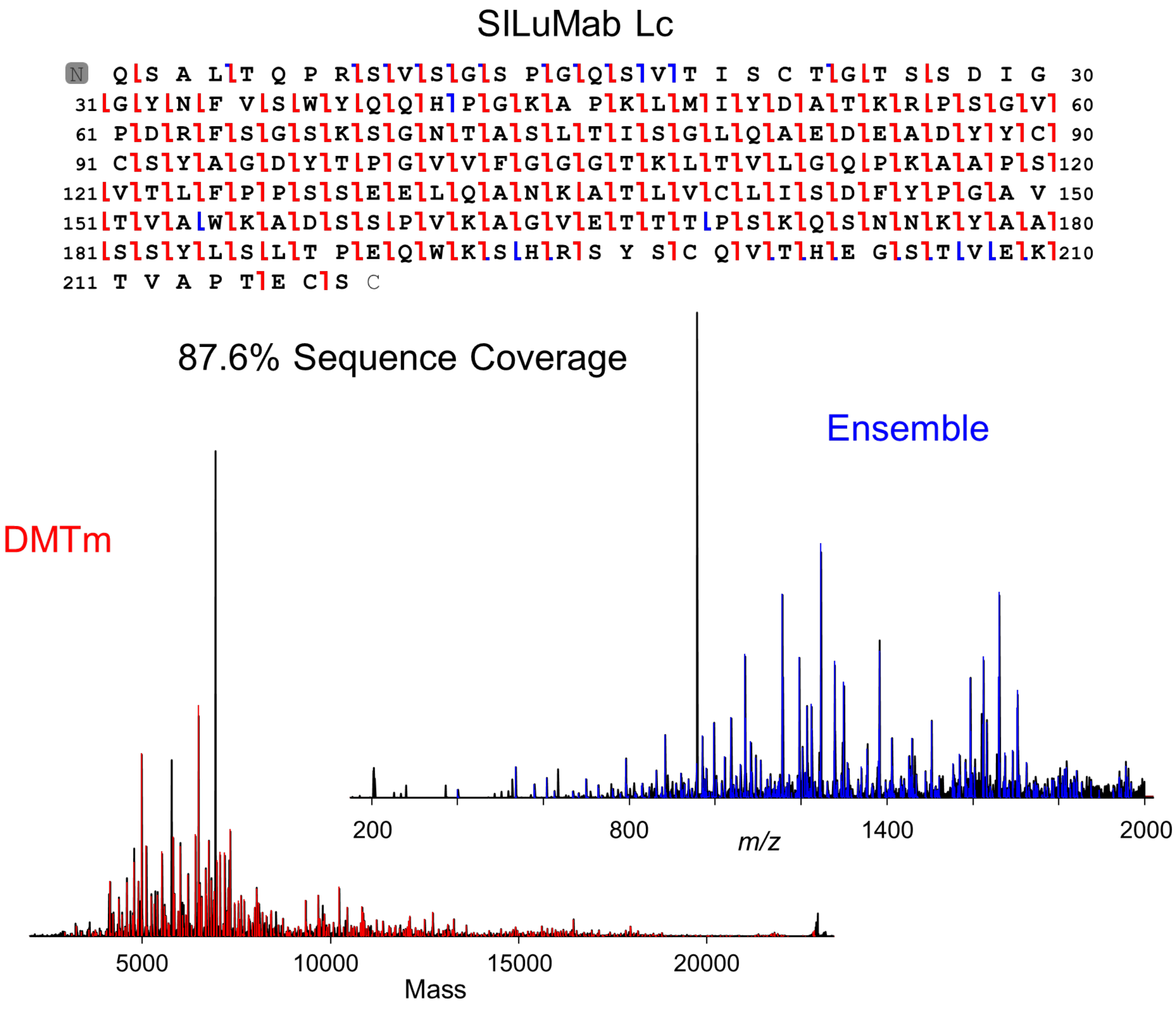


**Figure S12. Combined characterization of Lc subunit from SILuMab using DMTm and ensemble data**. The combined fragment map (top) is shown, and the annotated matching fragment ions are shown on the DMTm and ensemble spectra (bottom). Red and blue markers indicate ions identified by DMTm and ensemble mode, respectively.

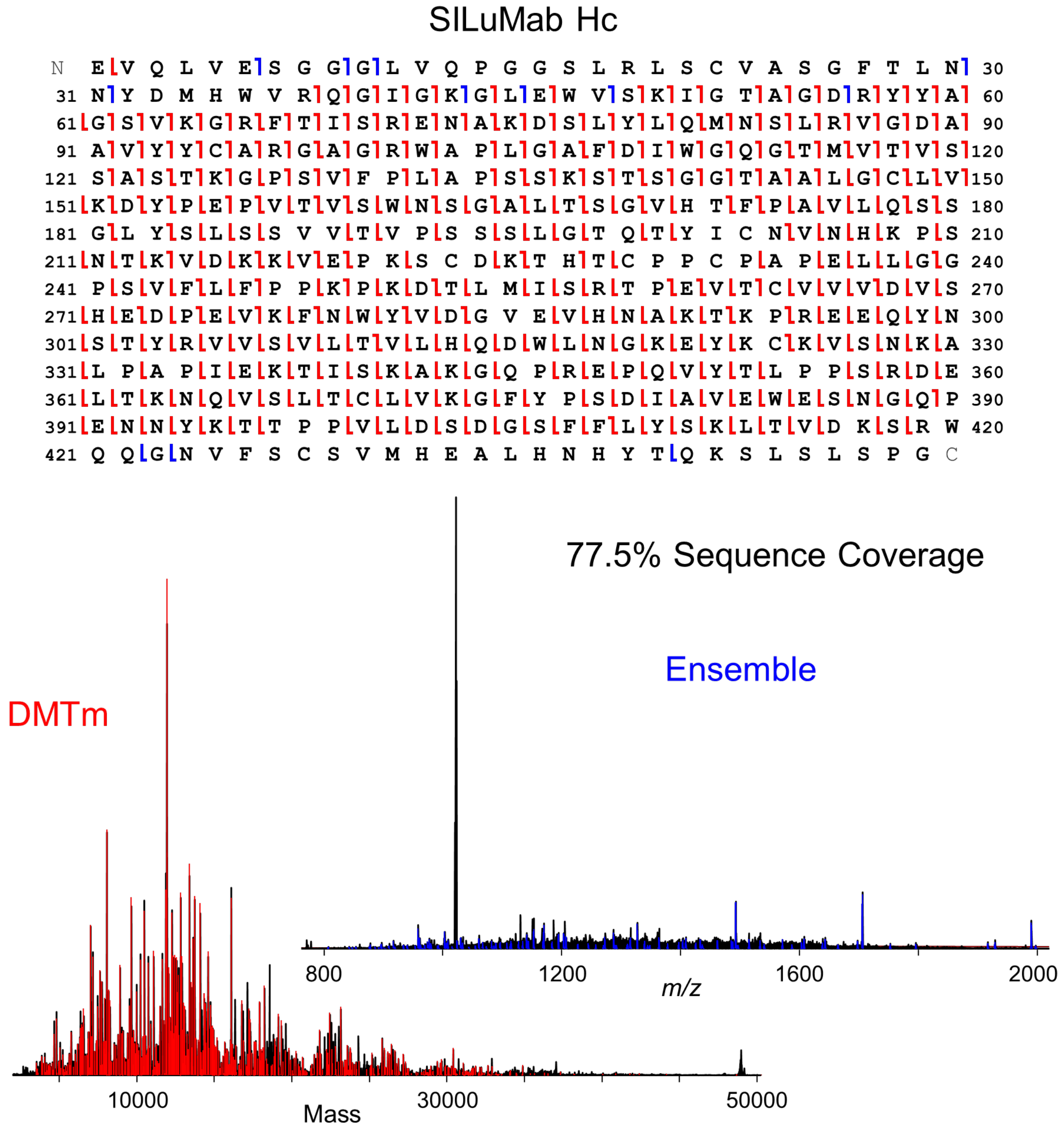


**Figure S13. Combined characterization of Hc subunit from SILuMab using DMTm and ensemble data**. The combined fragment map (top) is shown, and the annotated matching fragments are shown on the DMTm and ensemble spectra (bottom). Red and blue markers indicate ions identified by DMTm and ensemble mode, respectively.

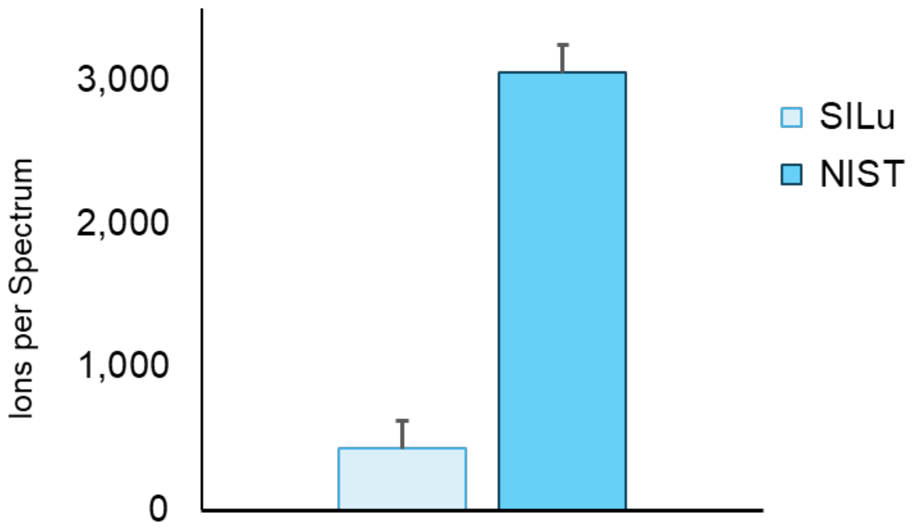


**Figure S14. Ion counts per spectrum.** The average counts of ions that were charge assigned per spectrum are shown for SILuMab using manual ion control and for NISTmab using automatic ion control (AIC).
